# HOXD13 is a direct EWS-FLI1 target and moderates fusion-dependent transcriptional states

**DOI:** 10.1101/2022.01.31.478548

**Authors:** April A. Apfelbaum, Feinan Wu, Brian Magnuson, Allegra G. Hawkins, Jennifer A. Jiménez, Sean D. Taylor, Elise R. Pfaltzgraff, Jane Y. Song, Cody Hall, Deneen M. Wellik, Mats Ljungman, Scott Furlan, Russell J.H. Ryan, Jay F. Sarthy, Elizabeth R. Lawlor

**Affiliations:** Cancer Biology PhD Program, University of Michigan, Ann Arbor, MI, 48109, USA; Seattle Children’s Research Institute, Seattle, WA, 98101, USA; Genomics and Bioinformatics Shared Resource, Fred Hutchinson Cancer Research Center, Seattle, WA, 98109, USA; Department of Biostatistics, University of Michigan, Ann Arbor, MI, 48109, USA; Childhood Cancer Data Lab Alex’s Lemonade Stand Foundation, Philadelphia, PA, USA; Department of Pediatrics, University of Michigan, Ann Arbor, MI, 48109, USA; Immunology Discovery, Genentech, Inc., 1 DNA Way, South San Francisco, CA 94080, USA; Department of Pathology, University of Michigan, Ann Arbor, MI, 48109, USA; Department of Cell and Regenerative Biology, University of Wisconsin, Madison, WI, 53705; Department of Radiation Oncology, University of Michigan, Ann Arbor, MI, 48109, USA; Fred Hutch Cancer Research Center, Seattle, WA, 98109, USA; Department of Pediatrics, University of Washington, Seattle, WA, 98105, USA

**Keywords:** EWS-FLI1, HOXD13, epigenetics, development, enhancer reprogramming, Ewing sarcoma, neuro-mesenchymal, cell state, CUT&RUN, CITE-seq

## Abstract

Oncogenic fusion proteins display exquisite tissue specificity, revealing that malignant transformation requires cooperation with cell-autonomous factors. Recent studies have also demonstrated that tumorigenicity of Ewing sarcoma requires precise regulation of the transcriptional activity of the EWS-FLI1 oncogenic driver. Here we show that the developmentally and anatomically restricted transcription factor *HOXD13* is a direct target of EWS-FLI1. Transcriptomic and CUT&RUN studies revealed that HOXD13 binds active, fusion-bound enhancers, resulting in altered expression of EWS-FLI1-induced targets. More strikingly, HOXD13 was found to bind and activate cis-regulatory regions of genes that are normally repressed by EWS-FLI1. Single-cell sequencing demonstrated marked intra-tumoral heterogeneity of HOXD13 transcriptional activity and revealed that antagonism between HOXD13-mediated gene activation and EWS-FLI1-dependent gene repression confers a spectrum of transcriptional cell states along a mesenchymal axis. Thus, HOXD13 serves as an internal rheostat for EWS-FLI1 activity, providing a paradigm for tissue-specific transcription factors as critical partners in fusion-driven cancers.

## Introduction

Oncogenic fusion proteins are common drivers of malignant tumors, especially in cancer affecting children and young adults (1). These fusions are often the sole recurrent mutation and individual fusions can be pathognomonic of a specific disease, demonstrating their exquisite tissue and context dependent nature (1). These observations, along with the recurring theme that oncogenic fusion proteins coopt normal developmental transcription programs, lead to the hypothesis that cooperation between oncogenic fusion proteins and developmental transcription factors may be essential for these fusions to exert their oncogenic effects.

Ewing sarcomas (EwS) are highly malignant, fusion-driven bone and soft tissue tumors that peak in adolescents and young adults (2). They are lethal cancers for over a third of diagnosed patients and for nearly all who develop metastatic disease (2). The tumors are highly undifferentiated histologically with phenotypic and ultrastructural features of both mesenchymal and neural lineages and are of presumed mesenchymal and/or neural crest stem cell origin (3–5). The genetic drivers of EwS arise from chromosomal translocations between a FET family member (*FUS/EWSR1/TAF15*) and an ETS family transcription factor, most commonly creating the EWS-FLI1 fusion (2). EWS-ETS proteins exert their oncogenic properties in large part *via* chromatin remodeling (6). Functioning as a pioneer factor, EWS-FLI1 creates *de novo* enhancers at GGAA microsatellite repeats throughout the genome, resulting in aberrant activation of normally heterochromatic regions and widespread transcriptional rewiring (6–10). In addition to its role in transcriptional activation, EWS-ETS fusions also lead to gene repression through not fully understood mechanisms (6, 11, 12).

Although EwS harbor few additional mutations (13, 14), they express a unique homeobox (HOX) gene profile that is distinct from other tumors and tissues (15–17). *HOX* transcription factors are critical for normal embryogenesis and their dysregulation can contribute to malignant transformation, as best exemplified by hijacking of HOXA9 in leukemogenesis (18, 19). In EwS, posterior *HOXD* genes (*HOXD10*, *HOXD11*, *HOXD13*) are highly expressed and HOXD13 contributes to tumorigenic and metastatic phenotypes (16, 17, 20). In normal development, HOXD13 contributes to limb development (21) and transcriptional regulation of mesenchymal gene programs (22, 23). The molecular mechanisms underlying HOXD13 activation and tumorigenic function in EwS have yet to be elucidated.

In the current work we show that HOXD13 is a direct epigenetic target of EWS-FLI1. In addition, our studies demonstrate that HOXD13 serves as a rheostat for EWS-FLI1 transcriptional activity wherein high levels of HOXD13 moderate expression of fusion-dependent target genes. Most strikingly, epigenomic and transcriptomic profiling data show that HOXD13 directly induces mesenchymal gene programs that are normally repressed by EWS-FLI1 and the relative activities of these two transcription factors determines EwS cell state along a mesenchymal axis. Thus, HOXD13 both cooperates with and competes with EWS-FLI1 in transcriptional regulation, providing a paradigm for tissue-specific transcription factors as critical partners in fusion-driven cancers.

## Materials & Methods

### Cell Culture

Ewing cell lines were obtained and cultured as previously described (24). H7-MSCs (kind gift of Dr. Sweet-Cordero) (25) were maintained in alpha-mem supplemented with 10% FBS, 2mmol/L-glutamine, and 1% Antibiotic-Antimycotic. Cells were cultured at 37°C with 5% CO2. Cells were all confirmed to be mycoplasma free and identities subject to STR-confirmation every 6 months.

### Lentivirus Production & Genetic Modification

Virus production and transductions carried out as previously described (24). pLKO.1 shNS and the shFLI1 were used for FLI1 knockdown experiments (Sigma, St. Louis, MO, USA). For HOXD13 knockdown experiments, stable cells lines were created with the doxycycline-inducible hairpins: pTripz shNS, shHOXD13 #1 or shHOXD13 #2. To induce the shRNA, 0.5 ug/mL doxycycline was added. EWS-FLI1 overexpression was achieved with pCLS-EGFP empty and EWS-FLI1-V5-2A-EGFP or pLIV empty and EWS-FLI1 (6).

### Quantitative PCR

Total RNA extraction, cDNA generation, and qPCR was performed as previously described (24). Primer and probe sequences are in Supplementary Table S1.

### Western Blot

Whole cell protein extraction, protein quantification, and western blot analysis was performed as previously described (24). Antibodies and the dilutions are in Supplementary Table S1. Membranes were imaged on the LiCor Odyssey imaging system.

### HOXD13 antibody production

Polyclonal anti-HOXD13 antibody was produced through peptide immunization in rabbits (YenZym, Brisbane, CA). The HOXD13 immunizing peptide described previously (23), was modified to contain a 16-amino-acid region with a Cysteine residue added to the N-terminus (C+VGLQQNALKSSPHASL) to facilitate coupling to a carrier protein.

### Immunofluorescence

Frozen sections of E13.5 *Hoxd13* WT, heterozygous, and knockout embryos (26) were formalin fixed, frozen in OCT and sectioned. Slides were thawed, washed, blocked, and incubated with the HOXD13 primary antibody in blocking buffer overnight (4C). Slides were incubated in donkey anti-rabbit 488 secondary for 1 hour (RT) followed by DAPI addition. Images were taken using an inverted Olympus IX83 (Tokyo, Japan) microscope with the CellSens Dimensions software. For immunocytochemistry, cells were fixed in 4% paraformaldehyde and permeabilized with 0.5% Triton. Cells were blocked for 1 hour (RT) and incubated either HOXD13 or IgG or 1 hour (RT). Following 3 washes, a fluorescent secondary antibody was added and incubated for 1 hour, followed by DAPI (1:10 000) incubation. Images were taken on a Lecia DMi8 microscope using the Lecia software.

### Migration assays

Real-time cell analysis (RTCA) of cell migration was performed as previously reported (27). 50 uL complete media was placed in the upper chambers and 160 uL serum-free media was added to the lower chambers before 1hr equilibration for at 37C. 5 x 10^4^ cells/well were plated in the upper chamber (100 uL) and plates were equilibrated for 30 minutes at room temperature. Migration was evaluated up to 30 hours.

### Mapping mouse enhancers to human

To map mouse enhancer sites in the regulatory HOXD domain the UCSC LIFTOVER tool was used to go from mm9 to hg19.

### Chromatin immunoprecipitation qPCR

Chromatin immunoprecipitation and qPCR was performed using the Zymo-Spin ChIP kit (Zymo, D5209) as previously described (24). Antibodies and primer sequences are in listed in Supplementary Table S1.

### CRISPRi two vector system

Cells transduced with UCOE-SFFV-KRAB-dCas9-P2A-mcherry were FACS sorted twice on mCherry expression. sgRNAs were designed flanking GGAA repeat sites using the Broad institutes GPP sgRNA designer. sgRNAs with high on-target scores were chosen and cloned into the sgOpti vector. Lentiviral sgRNA-containing sgOpti vectors were transduced into stably expressing dCas9-KRAB-mcherry cells and puromycin selected before collection after 8 days. Detailed protocol in Supplementary methods.

### Flow cytometry

Cells were washed with PBS + 2% FBS, followed by incubation with antibodies for 30 minutes at 4C in the dark. Cells were washed and analyzed on a BD accuri C6 machine. 10,000 events were collected, and live cells were gated on the unstained SSC vs FSC. To determine the percentage of positive cells, the isotype IgG control was used to set negative gates. Analyses were performed using FCS express (De Novo Software).

### Bru-seq/RNA-seq and analysis

Bru-seq was performed on A673 and CHLA10 cells following HOXD13 knockdown as described previously (28). Poly(A)-capture RNA-seq was performed for TC32 HOXD13 knockdown cells. Libraries were prepared with NEBNext Ultra II RNA Library Prep kit and paired end sequencing was performed on a Novaseq600. Reads were analyzed for quality control, trimmed, aligned to GRCh38, and analyzed for differential analysis (FastQC 0.1.9 (https://www.bioinformatics.babraham.ac.uk/projects/fastqc/), Trim Galore (Babraham Institute), STAR (29), and DESeq2 v1.18.1 (30)). Overrepresentation analysis was performed using the Broad Institute’s Molecular Signatures Database (MSigDB) (31). Heatmaps and volcano plots were made with the R packages pheatmap and EnhancedVolcano, respectively.

### Automated CUT&RUN Sequencing and analysis

Automated CUT&RUN protocol was performed as described (32) at the Fred Hutchinson Genomics core. Antibodies listed in Supplementary Table S1. FastQC 0.1.9 was used to examine read quality and paired-end reads were aligned to hg38 using Bowtie2.3.5 (33).

Narrow and broad peaks were called and filtered for histone marks using MACS2.2.7 (34). For HOXD13 peaks, both overlapping filtered MACS2 peaks and SEACR peaks (stringent mode) (35) were used. BEDTools 2.30.0 (36) was used to identify overlapping peaks between marks. Peak annotation and motif analysis was performed with HOMER 4.11 (37). The GeneOverlap and ChIPpeakAnno packages were used to calculate gene and genomic site overlap, respectively. Number of GGAA/CCTT sites were calculated for overlapped HOXD13 and EWS-FLI1 binding sites within 250bp up and downstream the peak in hg38. Detailed protocol in Supplementary methods.

### CITE-seq processing and analysis

CITE-seq allows for matched transcript and cell surface antigen profiling of individual cells (38, 39). 500 000 CHLA10 and A673 cells were resuspended in 25 uL Biolegend staining buffer (420201, San Diego, CA). 2.5 uL human TruStain FcX (422301, San Diego, CA) was added per sample (4C,10min). Hash-Tag Antibodies were incubated for 30 minutes (4C). Samples were pooled 1:1 at 1 000 cells/uL and libraries were generated using the 3’ V3 10X Genomics Chromium Controller (CG000183, Pleasanton, CA). Final library quality was assessed using the Tapestation 4200 (Agilent, Santa Clara, CA) and libraries were quantified by Kapa qPCR (Roche). Pooled libraries were subjected to paired-end sequencing on a NovaSeq 6000 (Illumina). Bcl2fastq2 Conversion Software (Illumina) was used to generate de-multiplexed Fastq files and the CellRanger (3.1) Pipeline (10X Genomics) was used to align reads and generate count matrices. Analysis performed using Seurat (40) and Monocle 3 (41, 42).

### Statistical analysis

Data were analyzed using GraphPad Prism software version 9.0 (San Diego, CA, USA). All statistical analyses were performed with Student’s t-test / One-way ANOVA followed by Tukey multiple comparison test / or Two-way ANOVA followed by Sidak’s multiple comparison test. Data are expressed as means and SEM from at least three independent experiments. Asterisk denoting *p*<0.05 (*) or *p*<0.01 (**).

## Data and Code Availability

Public data used in this study reported in Supplementary Table S1. Sequencing data generated in this study have been deposited at GEO (GSE182513). Code available at https://github.com/LawlorLab/HOXD13-Paper.

## Results

### HOXD13 expression in EwS is dependent on EWS-FLI1

Given that HOXD13 is uniquely highly expressed by EWS-ETS fusion-positive sarcomas (16), we hypothesized that *HOXD13* may be an EWS-FLI1 target. Knockdown of EWS-FLI1 in a panel of EwS cell lines (Fig. 1A-C, Supplementary Fig. S1A) led to downregulation of *HOXD13* with no appreciable change in expression of either *HOXD11* or *HOXD10* (Fig. 1D). Loss of HOXD13 protein was confirmed by immunocytochemistry (Fig. 1E) after successfully authenticating this custom antibody (Supplementary Fig. S1B-D). To validate the EWS-FLI1-dependent regulation of *HOXD13* in an orthogonal *in vivo* system, we interrogated published single-cell data from doxycycline-inducible EWS-FLI1 knockdown xenografts (43). Consistent with our *in vitro* studies, *HOXD13* expression was reduced upon loss of EWS-FLI1 (Fig. 1F). Thus, high levels of *HOXD13* in EwS cells are, at least in part, dependent on EWS-FLI1.

**Figure 1.**
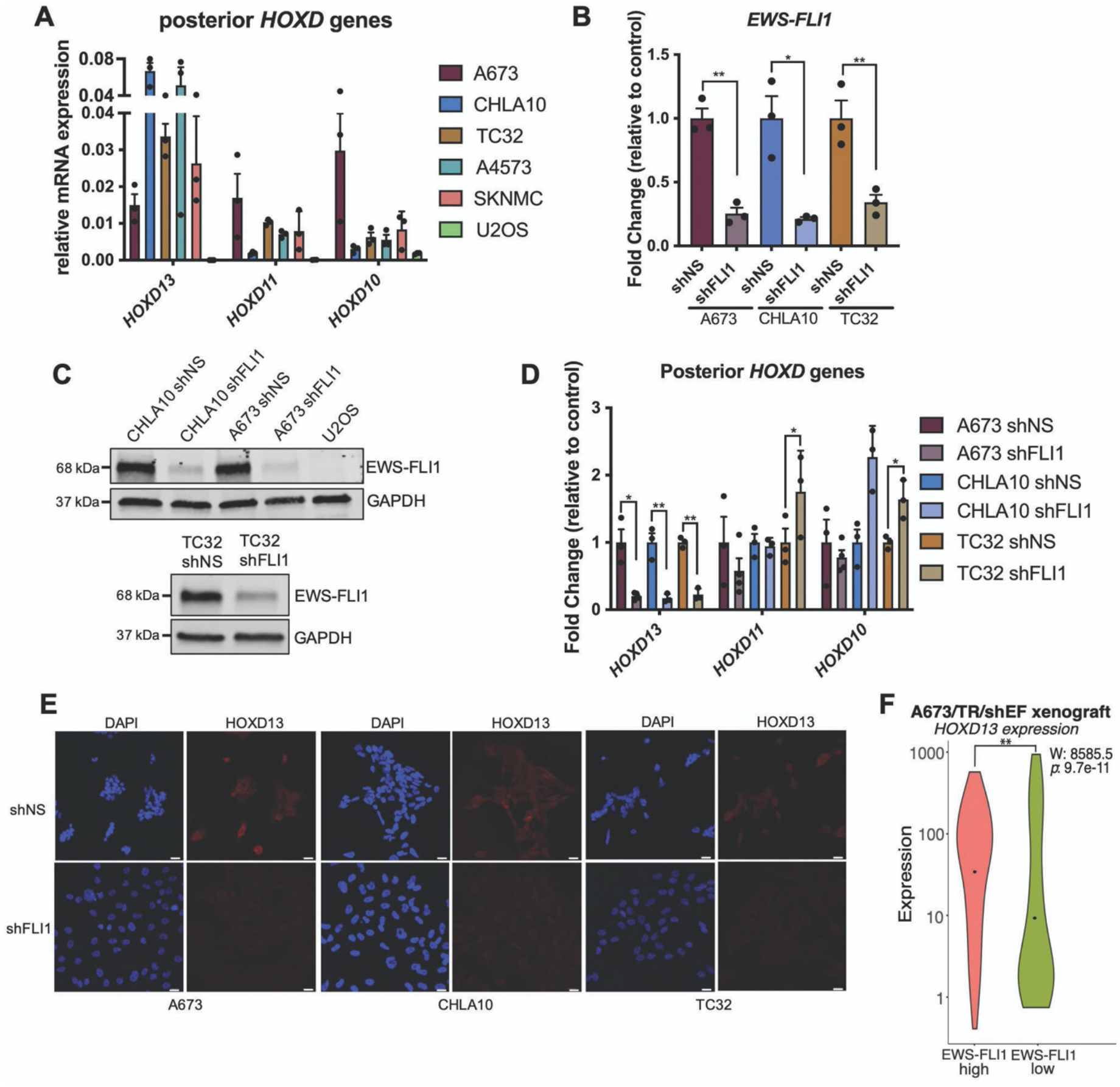
HOXD13 expression in EwS is dependent on EWS-FLI1. A) qRT-PCR of Posterior *HOXD* genes in EwS cells and U2OS (osteosarcoma) cells. B) qRT-PCR and C) western blot of EWS-FLI1 96 hrs after knockdown. D) qRT-PCR of *HOXD13*, *HOXD11*, and *HOXD10* expression in control and EWS-FLI1 knockdown cells. E) Fluorescent Immunocytochemistry of HOXD13 (Alexa647) in EWS-FLI1 knockdown cells. Nuclear counterstain was performed with DAPI. Scale bar is 10 um. F) Single-cell gene expression profiles from A673/TR/shEF xenografts (24) quantified by violin plots (Wilcoxon rank sum test). Error bars for qRT-PCR studies are representative of SEM from three independent replicates. Expression levels were determined relative to two housekeeping genes and fold changes expressed relative to control condition. * *p*<0.05; ** *p*<0.01; Two-way ANOVA; Sidak’s multiple comparison test; Two-tailed *t*-test.

### EWS-FLI1 creates and activates a *de novo HOXD13* enhancer in the developmentally conserved TAD

During embryogenesis, expression of HOX gene dosage and timing are tightly orchestrated by epigenetic mechanisms and expression of genes in the *HOXD* locus is specifically regulated by long-range enhancers in two developmentally conserved topologically associated domains (TADs) (44, 45). In murine development, *Hoxd13* is coordinated by five enhancers in a centromeric TAD (C-DOM) (Fig. 2A-Top) and disruption of this region leads to misexpression of *Hoxd13*, resulting in aberrant limb and posterior skeleton development (45). Given the established role of EWS-FLI1 as a pioneer factor (6, 9), we investigated whether the fusion might influence the chromatin state of this C-DOM region. To test this, we mapped the human C-DOM enhancers from their syntenic regions in mice using the UCSC LiftOver tool (Fig. 2A). The C-DOM region in mice, and corresponding syntenic region in humans, is a 600kb gene desert that starts approximately 180kb upstream (5’) of the *Hoxd13* promoter (45). We interrogated this region for potential EWS-FLI1 binding sites and identified a 14-consecutive repeat GGAA microsatellite (Fig. 2A). Published chromatin immunoprecipitation (ChIP)-seq data from EwS cells and tumors (6) confirmed EWS-FLI1 binding and H3K27ac and H3K4me1 histone modifications at this site (Fig. 2A & B). In keeping with EWS-FLI1-dependent enhancer regulation, the activating histone marks were lost upon EWS-FLI1 knockdown (Fig. 2C) and non-Ewing sarcoma tumors (46) showed no evidence of chromatin activation (Fig 2B).

**Figure 2.**
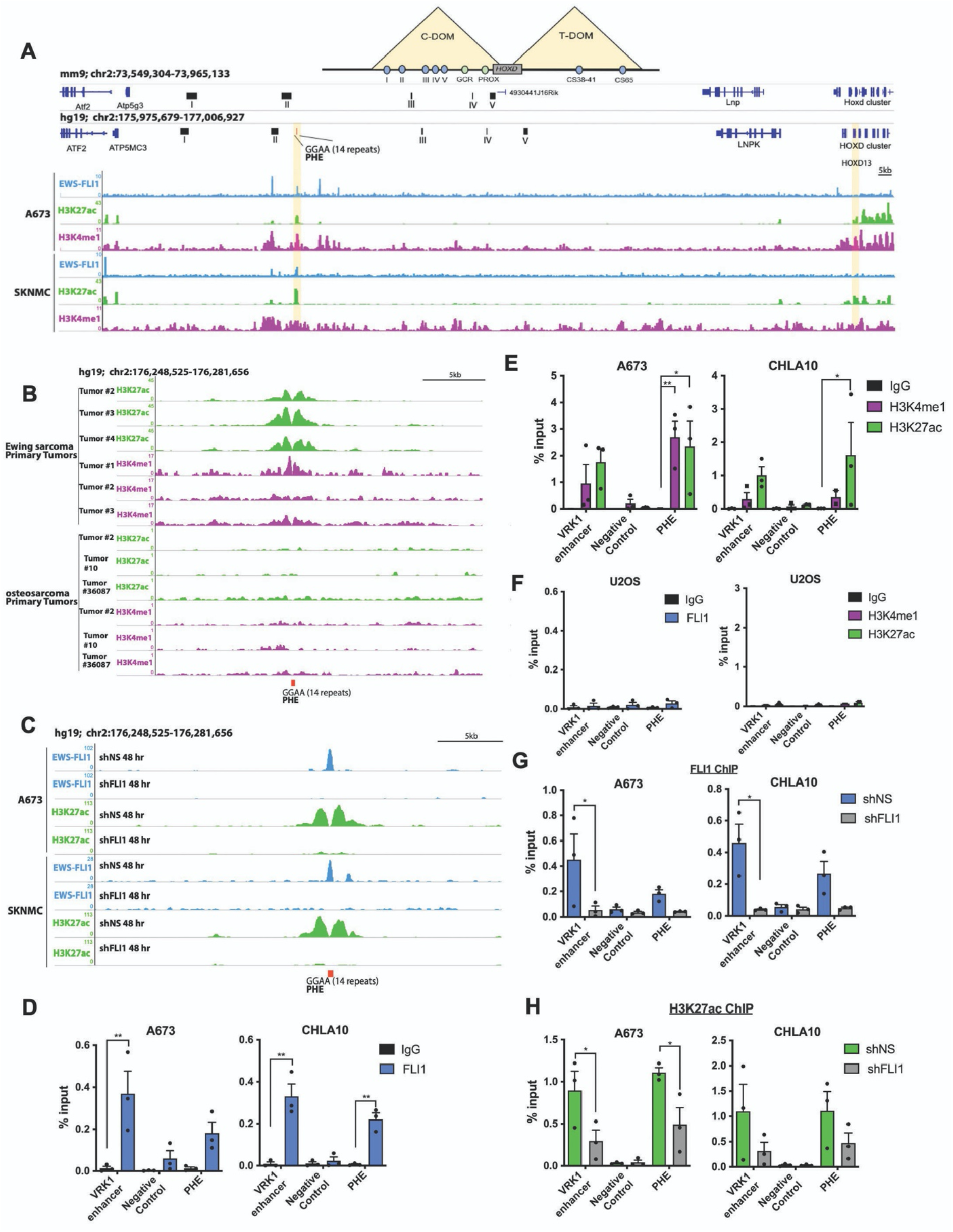
EWS-FLI1 creates and activates a *de novo HOXD13* enhancer in the developmentally conserved TAD. A) The murine HOXD C-DOM region (mm9) and its corresponding syntenic region in human (hg19) with annotation of human-specific GGAA microsatellite site (posterior HOX enhancer: PHE). ChIP-seq tracks of EWS-FLI1, H3K27ac, and H3K4me1 binding at the PHE region in EwS cells (6). B) ChIP-seq tracks of H3K27ac and H3K4me1 at the PHE of primary EwS (6) and osteosarcoma (28) tumor samples. C) ChIP-seq tracks of EWS-FLI1 and H3K27ac at the PHE region in EwS cells following EWS-FLI1 knockdown (6). D) ChIP-qPCR for EWS-FLI1, E) H3K27ac, and H3K4me1 in EwS cells. F) ChIP-qPCR for EWS-FLI1, H3K27ac, and H3K4me1 in U2OS cells. ChIP-qPCR for G) EWS-FLI1, H) H3K27ac after EWS-FLI1 knockdown. Negative control: an inert intergenic region in chr2. VRK1 enhancer: positive control GGAA enhancer site (7). C-DOM: centromeric domain; T-DOM: telomeric domain. Error bars representative of SEM from three independent replicates. * *p*<0.05; ** *p*<0.01; Two-way ANOVA; Sidak’s multiple comparison test; Two-tailed *t*-test.

We next sought to directly test and validate this GGAA microsatellite, named the <u>p</u>osterior <u>H</u>OXD <u>e</u>nhancer (PHE), as an EWS-FLI1-dependent enhancer through ChIP-qPCR (6, 9). With VRK1 enhancer as a positive control, these studies confirmed EWS-FLI1 binding and H3K27ac/H3K4me1 marks at the PHE (Fig. 2D and 2E) in EwS cells but not in U2OS osteosarcoma cells (Fig. 2F). Knockdown of EWS-FLI1 led to loss of fusion-binding (Fig. 2G) and concomitant loss of H3K27ac enrichment (Fig. 2H). Thus, in EwS cells, EWS-FLI1 binds and activates a *de novo* GGAA microsatellite enhancer in the HOXD C-DOM TAD regulatory domain.

### The PHE uniquely controls *HOXD13* expression in Ewing sarcoma

To functionally validate the PHE as an enhancer, we used CRISPR interference (CRISPR-dCas9-KRAB; CRISPRi) to focally induce a H3K9me3-marked repressive chromatin state (47, 48). Since GGAA sites are repetitive and non-specific, we designed three unique sgRNAs that flank the PHE. In parallel to PHE-targeted sgRNAs, dCas9-KRAB expressing cells were transduced with validated control sgRNAs that target either the SOX2 GGAA enhancer (10) or a non-coding, inert genomic region (49). Only cells that were transduced with PHE-targeting sgRNAs acquired the H3K9me3 mark at the PHE locus (Fig. 3A). Similarly, cells transduced with the SOX2 GGAA-targeted sgRNA acquired the H3K9me3 mark at the SOX2 enhancer (Supplementary Fig. S2A). Conversely, SOX2 GGAA-targeted sgRNA had no impact on the chromatin state of the PHE and *vice versa* (Fig. 3A; Supplementary Fig. S2A).

**Figure 3.**
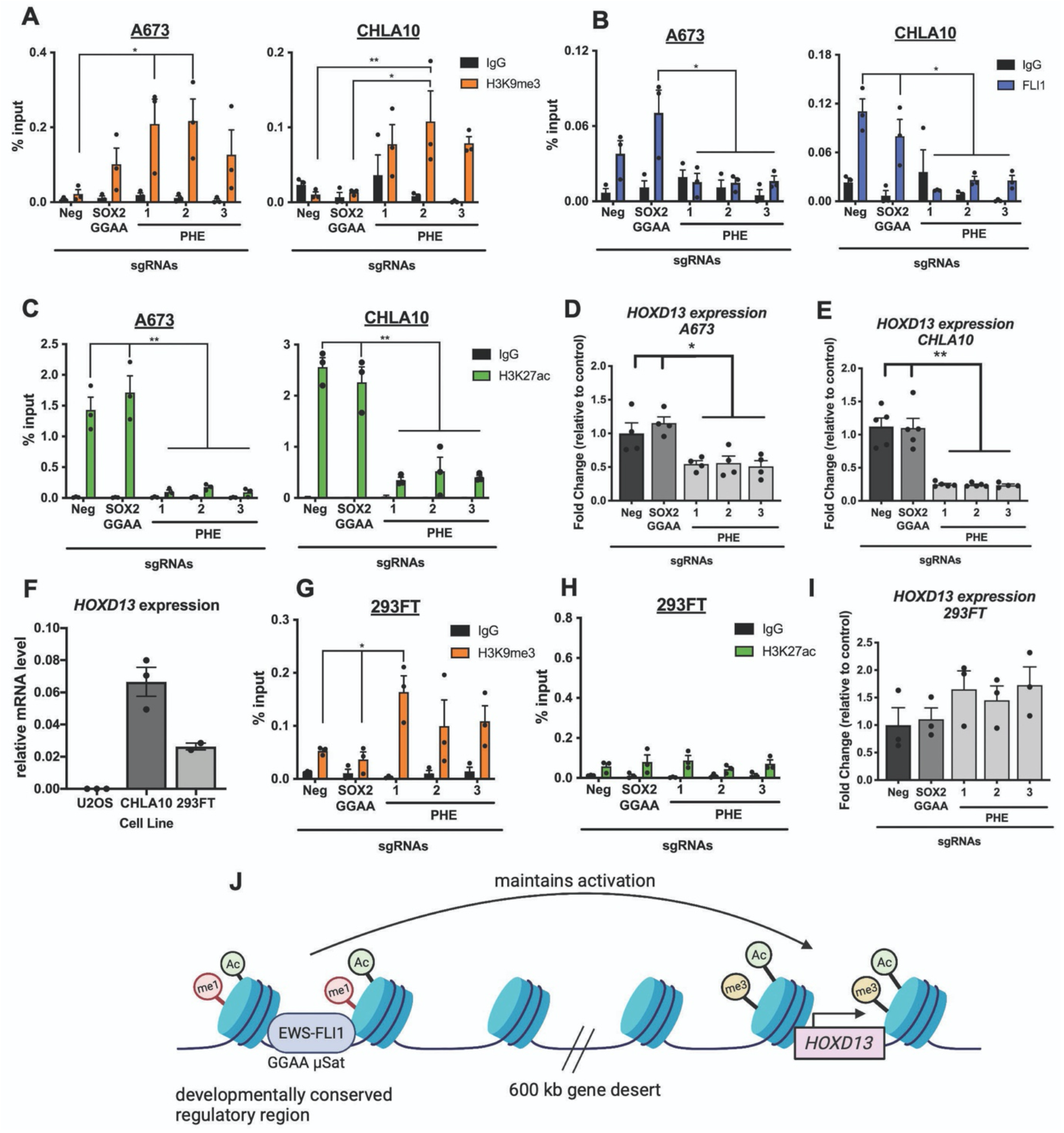
The PHE uniquely controls *HOXD13* expression in Ewing sarcoma. All cells express dCas9-KRAB-mcherry and sgRNAs targeting either a negative control region (neg), the SOX2 GGAA enhancer, or PHE (1–3). DNA was isolated 8 days after sgRNA transduction. A) ChIP-qPCR for H3K9me3, B) EWS-FLI1, and C) H3K27ac at the PHE region. qRT-PCR of *HOXD13* levels in D) A673 and E) CHLA10 cells. F) qRT-PCR of baseline *HOXD13* expression in U20S, CHLA10, and 293FT cells. ChIP-qPCR for G) H3K9me3 and H) H3K27ac at the PHE region in gRNA-targeted 293FT cells. I) qRT-qPCR of *HOXD13* expression in PHE-targeted 293FT cells. J) Model of EWS-FLI1 regulation of HOXD13 in Ewing sarcoma cells (biorender). Error bars are representative of SEM from at least three independent experiments. * *p*<0.05; ** *p*<0.01; Two tailed t-test; Two-way ANOVA; Sidak’s multiple comparison test.

We next evaluated the impact of H3K9me3 deposition on EWS-FLI1 binding and enhancer activation. Acquisition of H3K9me3 at the PHE resulted in site-specific loss of EWS-FLI1 binding (Fig. 3B). Further, targeted gain of H3K9me3 at the PHE was accompanied by a striking loss of H3K27ac at the targeted locus (Fig. 3C & Supplementary Fig. S2B-D) and by reduced *HOXD13* expression (Fig. 3D-E). Since 293FT cells also express *HOXD13* (Fig. 3F) in the absence of an EWS-ETS fusion, we assessed the effect of targeting the PHE in these cells. Induction of heterochromatin at the PHE in 293FT cells affected neither H3K27ac nor *HOXD13* expression (Fig. 3G-I). Thus, the EWS-FLI1-bound PHE GGAA microsatellite functions as a distal enhancer in EwS, contributing to transcriptional activation of *HOXD13* (Fig. 3J).

Analysis of published ATAC-seq and ChIP-seq data from human MSCs (6, 9) showed that, when expressed in these cells of putative tumor origin, EWS-FLI1 binds to the PHE and induces an open and active chromatin state (Supplementary Fig. S3D-E). Nevertheless, despite this clear pattern of chromatin remodeling, *HOXD13* transcription was not induced in these cells (6, 9). Consistent with this, we also did not detect any upregulation of *HOXD13* expression in human MSCs, U2OS osteosarcoma, or SW1353 chondrosarcoma cells following ectopic expression of EWS-FLI1 (Supplementary Fig. S3A-C). Thus, although direct EWS-FLI1 binding of the PHE reproducibly leads to epigenetic rewiring and creation of a *de novo* enhancer element, this enhancer hijacking is, by itself, insufficient to induce *HOXD13* expression (Fig. 3J).

### HOXD13 regulates mesenchymal gene programs and cell states

To elucidate how HOXD13 effects its oncogenic function in EwS, we performed RNA-seq on three independent human tumor-derived EwS cell lines following doxycycline-induced knockdown of *HOXD13* (Fig. 4A). Highly significant and robust changes in transcriptomes were universally observed demonstrating that HOXD13 is highly transcriptionally active in EwS cells (Supplementary Fig. S4A-C, Supplementary Table S2). However, the specific identity of HOXD13-regulated gene targets was highly cell line-dependent revealing the importance of cell context for HOXD13 activity. To identify the gene programs that are most likely to be relevant for EwS phenotypes we focused on HOXD13-regulated genes that were common to all three cell lines (Fig. 4B). Of 119 shared genes, 109 were regulated in the same direction and most (N=87) were downregulated, indicating that they are positively regulated by HOXD13 (Fig. 4C). Analysis of gene ontologies revealed enrichment of mesenchymal programs among these HOXD13-activated targets (i.e. downregulated upon loss of HOXD13), while neural differentiation and development genes were more prominent among transcripts that are suppressed by HOXD13 (Fig. 4D-E; Supplementary Table S3). Notably, transcription factors and markers of neural and mesenchymal differentiation and genes involved in epithelial mesenchymal transitions (EMT) and metastasis phenotypes were prominent among HOXD13-regulated transcripts (Fig. 4C).

**Figure 4.**
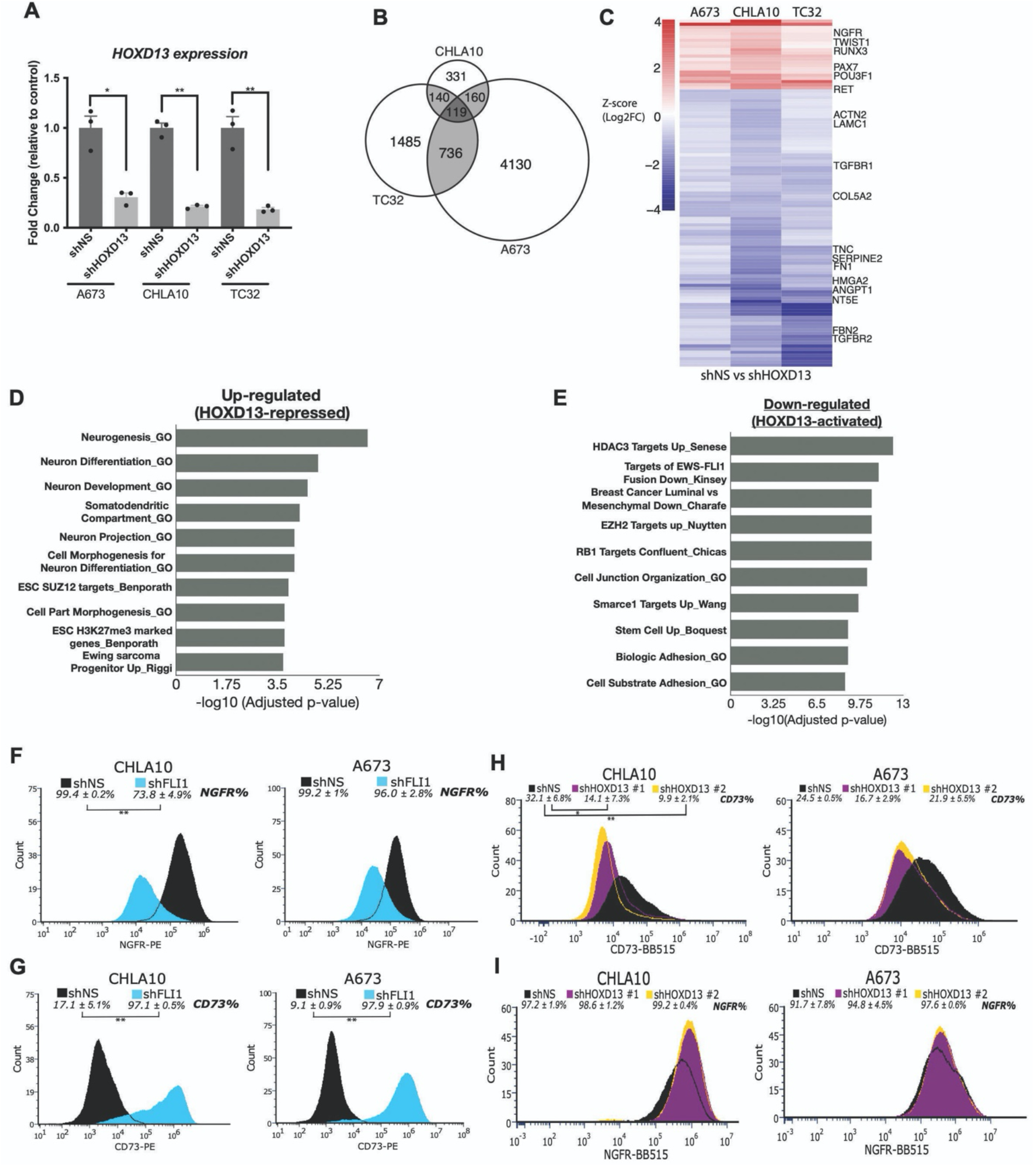
HOXD13 regulates neuro-mesenchymal gene programs and influences cell states. A) qRT-PCR of *HOXD13* expression in cells submitted for RNAseq. B) Venn diagram showing overlap of significantly differentially expressed genes in each cell line. C) Heatmap depicting the differentially expressed genes (*padj <0.05*) between shHOXD13 and shNS for all cell lines. Scale: Z-score (Log2FC). Overrepresentation analysis of top 10 gene sets D) up- and E) down-regulated following HOXD13 knockdown in all cell lines. Flow cytometry histograms showing the shift in F) NGFR + cells and G) CD73+ cells upon EWS-FLI1 knockdown. Flow cytometry histograms showing the shift in H) NGFR + cells and I) CD73+ cells upon HOXD13 knockdown. Flow Error represents SD from at least three independent experiments. * *p*<0.05; ** *p*<0.01; Two-tailed *t*-test.

EwS tumors display features of both neural and mesenchymal lineages and EWS-FLI1 has been shown to promote neural-, whilst inhibiting mesenchymal-like states (3, 50) (51). In view of our transcriptomic results, we hypothesized that the relative mesenchymal state of EwS may be in part controlled by HOXD13. The MSC marker *NT5E* (ecto-5’-nucleotidase; CD73) was among HOXD13-induced transcripts and *NGFR*, a neural marker, was relatively repressed by HOXD13 (Fig. 4C). Thus, we used these cell surface proteins to mark neural-like (NGFR+) and mesenchymal-like (CD73+) EwS cells in control and genetically modified conditions. As shown, most EwS cells express NGFR and, as expected from prior literature (3, 50) (51), knockdown of EWS-FLI1 resulted in loss of NGFR+ cells and gain in the frequency of CD73+ cells (Fig. 4F-G). In direct contrast, HOXD13 knockdown led to more robust cell surface NGFR expression and to diminished numbers of mesenchymal-like CD73+ cells (Fig. 4H-I). In addition, HOXD13 knockdown cells were reproducibly less migratory than their respective controls (Supplementary Fig. S4D-F). Thus, HOXD13 promotes mesenchymal gene programs and phenotypes in EwS.

### HOXD13 binds active chromatin in EwS cells at intergenic and intronic sites and at EWS-ETS binding sites

The transcriptional profiling of HOXD13 knockdown cells revealed that numerous genes that are positively regulated by HOXD13 are normally repressed by EWS-FLI1 (Fig. 4E), while EWS-FLI1-induced genes are over-represented among genes HOXD13-repressed genes (Fig. 4D). This antagonistic effect of HOXD13 on EWS-FLI1-dependent gene regulation was not due to changes in the level of the fusion (Supplemental Fig. S4G-J). To determine how HOXD13 influences EWS-FLI1-dependent transcriptional activity, we performed CUT&RUN-sequencing to identify HOXD13 binding sites in EwS cells and define chromatin states at these sites (52). Binding of HOXD13 protein was detected at thousands of sites throughout the genome in both cell lines, with nearly 500 binding sites shared at predominantly intronic and intergenic regions (Fig. 5A; Supplementary Fig. S5A, Supplementary Table S4). CUT&RUN-sequencing showed enrichment of H3K27ac and H3K4me1 at these HOXD13 binding sites, identifying them as putative active enhancers (Fig. 5B-C, Supplementary Fig. S5B-C). Likewise, HOXD13-bound promoter/transcription start sites were characterized by enrichment of active chromatin marks, H3K27ac and H3K4me3 (Fig. 5B-C, Supplementary Fig. S5B-C). No enrichment of the repressive H3K27me3 mark was detected at any HOXD13 bound sites (Fig. 5B-C, Supplementary Fig. S5B-C).

**Figure 5.**
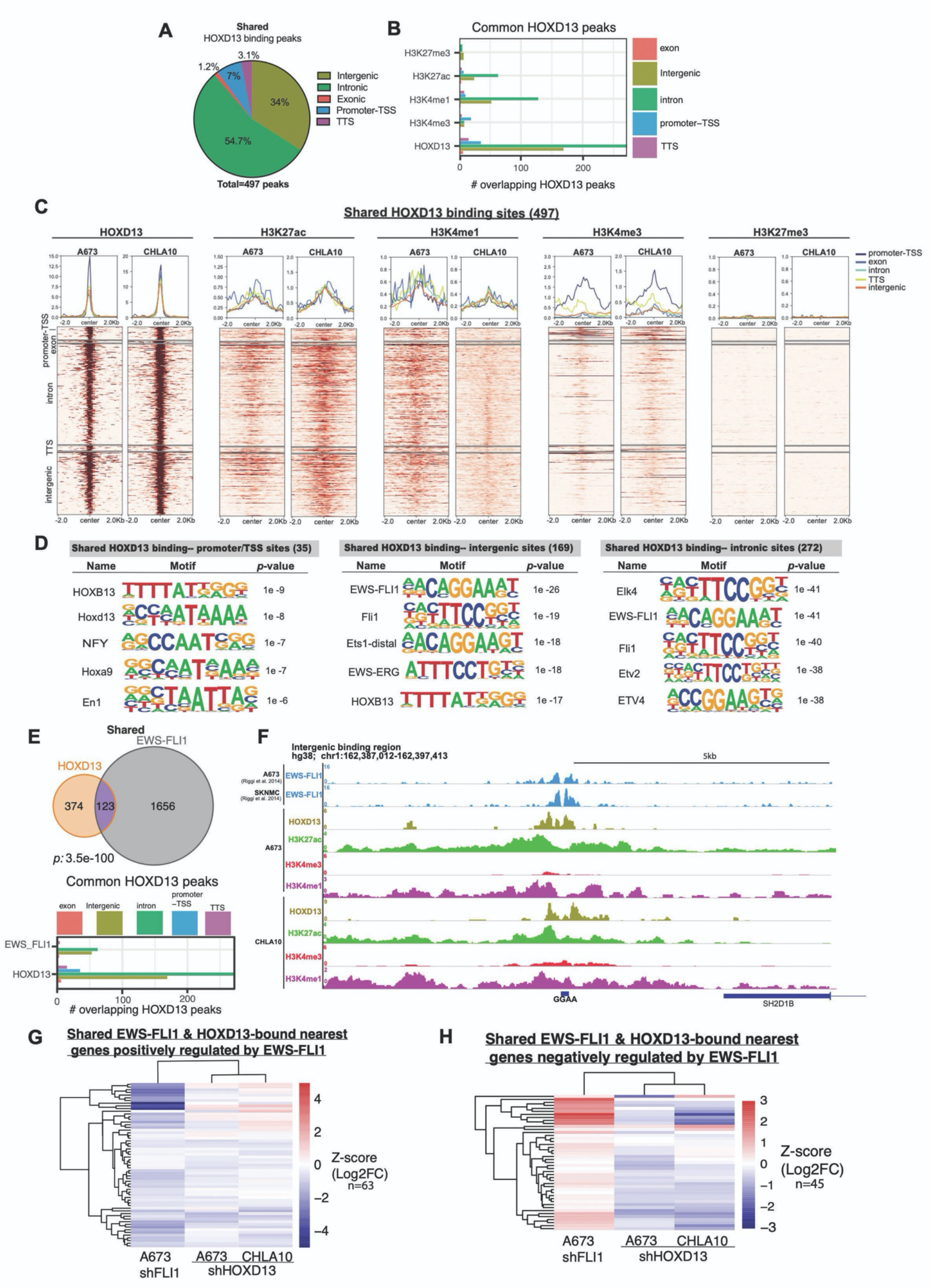
HOXD13 binds active chromatin in EwS cells at intergenic and intronic regions and at EWS-ETS binding sites. A) Pie chart showing the genomic distribution of HOXD13 binding sites shared between CHLA10 and A673 cells. B) Bar chart summarizing shared binding sites and associated histone marks at these sites. C) Tornado plots depicting shared HOXD13 binding and the associated histone marks by genomic location. D) HOMER Motif analysis by genomic location for the shared HOXD13 binding sites. E) Venn diagrams showing the overlap between HOXD13 binding sites and published EWS-FLI1 binding sites in shared sites. Bar graphs depict genomic locations of shared HOXD13 and EWS-FLI1 bound sites. F) Representative CUT&RUN tracks of HOXD13 binding and associated histone marks at a direct target intronic region. G) Heatmaps depicting shared HOXD13 and EWS-FLI1 nearest genes negatively regulated by EWS-FLI1 after HOXD13 knockdown. H) Heatmaps depicting shared HOXD13 and EWS-FLI1 nearest genes negatively regulated by EWS-FLI1 after HOXD13 knockdown.

In normal development, HOX proteins have DNA binding affinities that are determined by their interactions with cell context-dependent cofactors (53, 54). Therefore, we reasoned that identification of enriched transcription factor binding motifs at HOXD13-bound loci would provide insights into its regulatory partners in EwS. HOMER analysis revealed the expected enrichment of HOX and other early developmental transcription factor motifs at gene promoter/transcription start sites (Fig. 5D, Supplementary Fig. S5D-E, Supplementary Table S5). In contrast, intergenic and intronic peaks showed a striking and reproducible enrichment of ETS family binding motifs, including EWS-ETS sites (Fig. 5D, Supplementary Fig. S5D-E). To directly test whether HOXD13 peaks localized to EWS-ETS binding sites, we compared HOXD13 peaks in A673 and CHLA10 cells to nearly 1 800 sites that were previously identified as EWS-FLI1-bound regions in A673 and SKNMC cells (6). As shown, a striking and highly statistically significant overlap exists between HOXD13 and EWS-FLI1-bound sites, especially in intergenic and intronic regions (Fig. 5E, Supplementary Fig. S5F-G, Supplementary Table S6). Although just over half of these shared peaks occur at GGAA repeats, the remainder do not, demonstrating that shared loci are not defined by the presence or absence of microsatellites (Fig. 5F, Supplementary Fig. S5H).

To determine the impact of HOXD13 at fusion-bound loci, we mapped the 123 shared binding sites to their 108 nearest genes as previously described (6), and assessed how modulation of either transcription factor influenced gene expression. Consistent with the established role of EWS-FLI1 as both an activator and repressor of gene transcription, approximately two-thirds of co-bound loci (63/108, log2FC <0) are positively regulated by the fusion (Fig. 5G), while the remainder (45/108, log2FC >0) are relatively repressed (Fig. 5H). In contrast, nearly all shared loci are activated by HOXD13 (Fig. 5G & H). Thus, binding of HOXD13 at EWS-FLI1-bound loci promotes activation of target genes, irrespective of how the gene is regulated by the fusion. As such, HOXD13 binding can augment expression of direct EWS-FLI1 target genes that are normally induced by the fusion and activate genes that are normally subject to EWS-FLI1-mediated silencing.

### Transcriptional antagonism between HOXD13 and EWS-FLI1 is largely indirect and evident at single-cell resolution

Although a subset of HOXD13 binding sites are established EWS-FLI1-bound loci, most are not. We therefore sought to broadly define direct transcriptional targets of HOXD13 in EwS cells. Using a nearest gene approach and integration of RNAseq and CUT&RUN data, we identified genes that were both bound and regulated by HOXD13 in A673 and CHLA10 cells (Fig. 6A-B). Consistent with its overall distribution, HOXD13 binding sites were present in adjacent introns or upstream intergenic regions of its direct target genes (Fig. 6C-D). In addition, most direct target genes were found to be activated by HOXD13 (Fig. 6E-F). Significantly, GSEA of directly activated target genes in both cell lines revealed enrichment of the EWS-FLI1-repressed signature (Fig. 6G-H). Thus, in EwS cells, HOXD13 directly binds and activates cis-regulatory regions of its transcriptional targets and many of these HOXD13-activated genes are genes that are normally repressed, either directly or indirectly, by EWS-FLI1.

**Figure 6.**
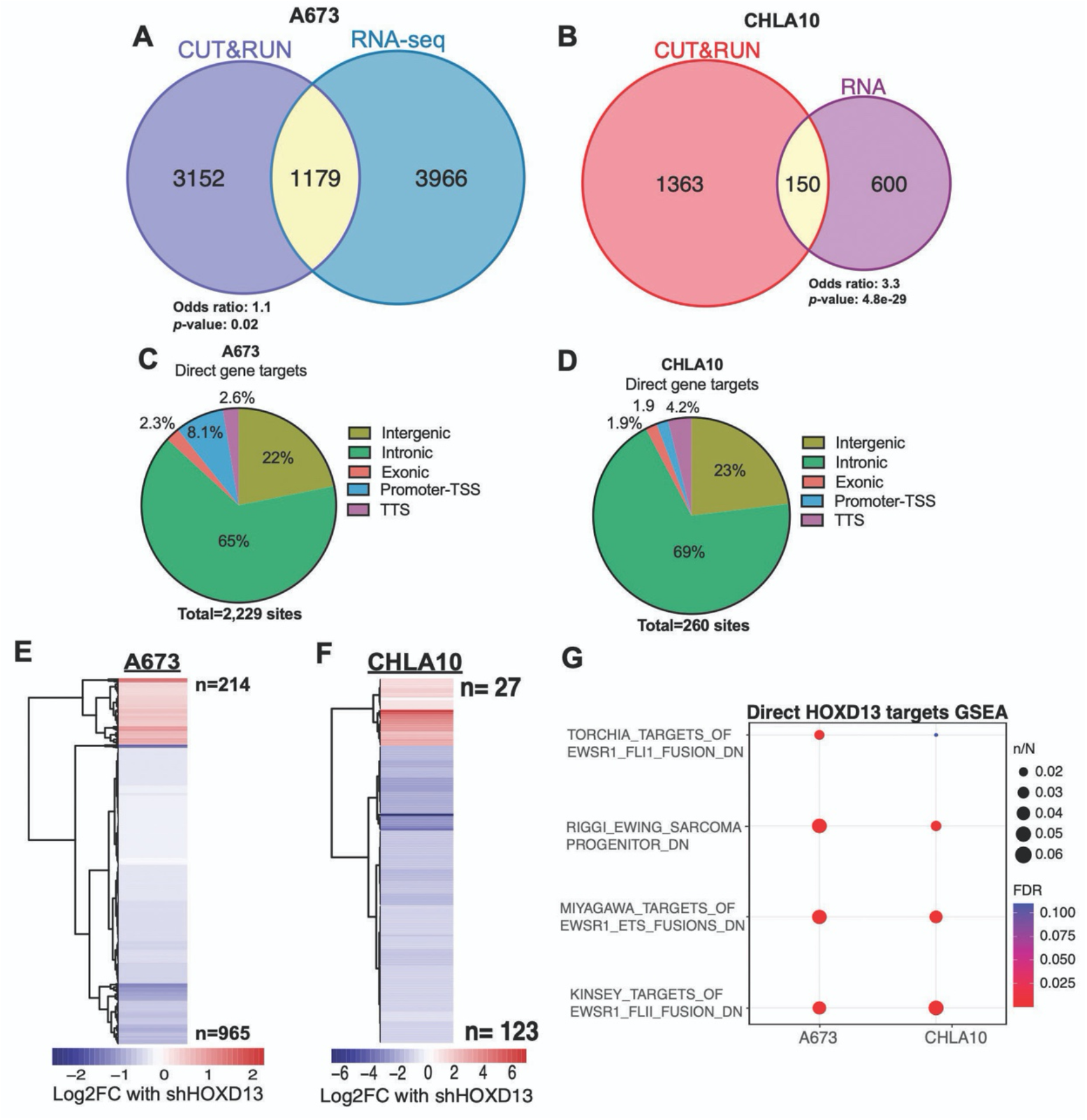
HOXD13 directly activates EWS-FLI1 repressed genes. A-B) Venn diagrams of the overlap between HOXD13 bound (CUT&RUN) and regulated (RNA-seq) genes in each cell line. C-D) Pie charts depict the genomic distribution of these sites. E-F) Heatmaps show the relative change in expression of these “direct” targets with HOXD13 knockdown. G) Overrepresentation analysis of direct HOXD13 target genes.

These studies of bulk populations established that transcriptional antagonism exists between HOXD13 and EWS-FLI1 but could not elucidate whether this antagonism exists at the level of individual tumor cells. To address this, we performed single cell-sequencing of A673 and CHLA10 cells using Cellular Indexing of Transcriptomes and Epitopes by Sequencing (CITE-seq) (39)(Supplementary Fig. S6A). The results showed that *HOXD13* expression is highly heterogeneous, both within and between cell lines (Fig. 7A). Similarly, and consistent with prior reports (43, 55), inter- and intra-cell line heterogeneity of EWS-FLI1 is evident, though variability is considerably less than *HOXD13* (Fig. 7A). Of note, given that direct quantification of the fusion transcript is not feasible using short read sequencing methods, we used the recently published EWS-FLI1-specific signature (IC-EwS) (43) as a surrogate to infer EWS-FLI1 expression. No correlation was detected between expression of *HOXD13* or the IC-EwS signature at the level of individual cells suggesting that transcriptional antagonism between HOXD13 and EWS-FLI1 cannot be fully explained by differences in the absolute levels of each gene (Fig. 7B). Next, we quantified the relative transcriptional activities of each transcription factor in individual cells. Transcriptional activity of the fusion in individual cells was determined by quantifying the relative expression level of established activated (N=1 244) and repressed (N=319) EWS-FLI1 target gene signatures (56). The HOXD13 transcriptional activation signature was derived from CHLA10 and A673 cells (N=254 genes). At the level of individual cell transcriptomes, no correlation was detected between HOXD13 activity and expression of EWS-FLI1-activated genes (Fig. 7C, Supplementary Fig. S6B&C). In contrast, a significant and reproducible direct correlation was observed between HOXD13 activity and expression of EWS-FLI1-repressed genes (Fig. 7D, Supplementary Fig. S6B&D). Moreover, this pattern was also evident in published single-cell data generated from five patient derived-xenografts (PDX) (43) (Fig. 7E-G, Supplementary Fig. S6E-F). Thus, the direct relationship between HOXD13-activation and upregulated expression of EWS-FLI1-repressed genes is evident in individual EwS cells both *in vitro* and *in vivo*.

**Figure 7.**
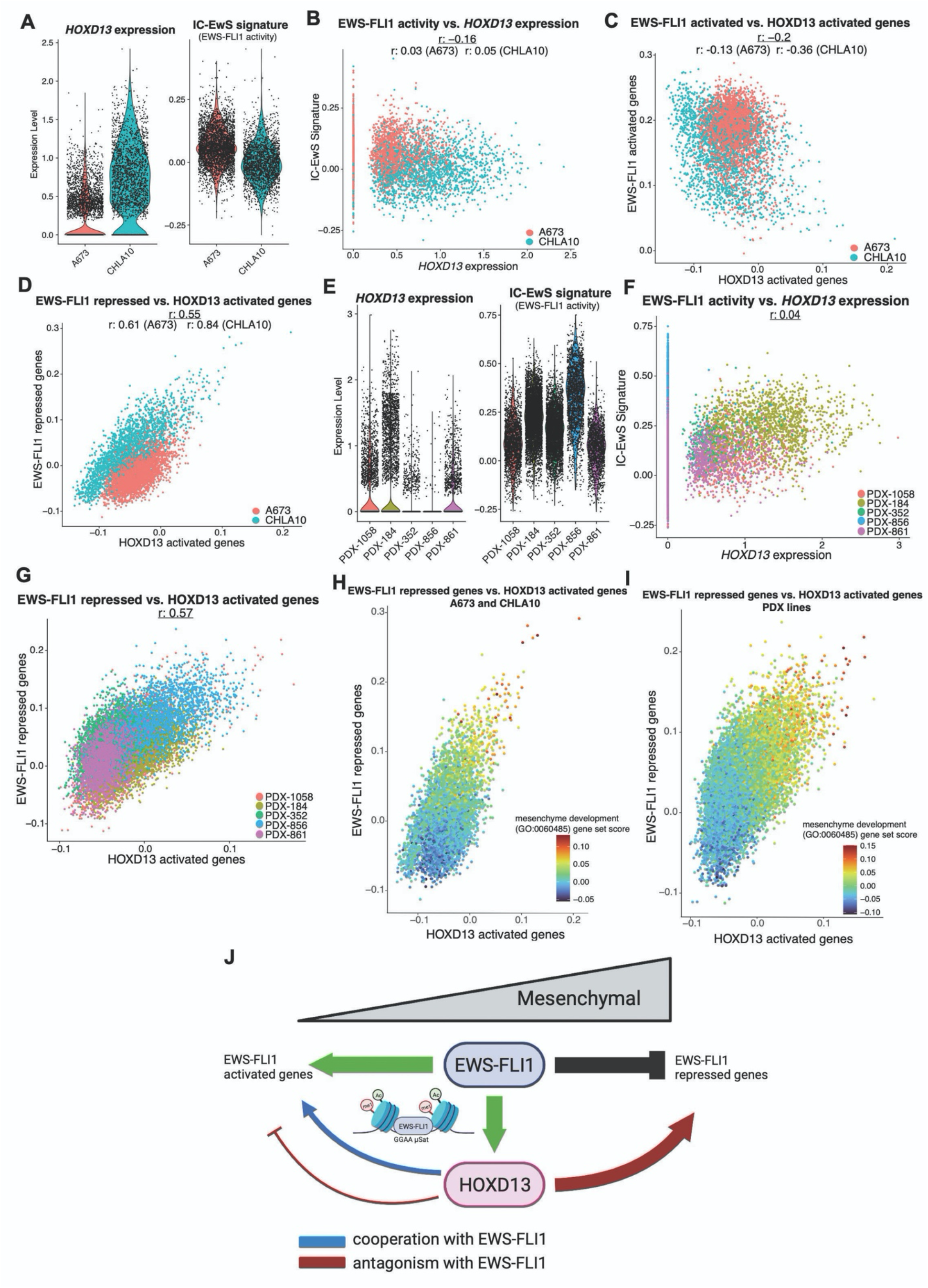
Transcriptional antagonism between HOXD13 and EWS-FLI1 activity is largely indirect and evident at single-cell resolution. Data were generated using CITE-seq. **A)** Violin plots showing single-cell HOXD13 expression and the IC-EwS EWS-FLI1 signature in A673 and CHLA10 cells. **B)** Scatter plot of *HOXD13* expression and the IC-EwS EWS-FLI1 signature by cell line. **C)** Scatter plot of HOXD13 activated genes and EWS-FLI1 activated genes by cell line. **D)** Scatter plot of HOXD13 activated genes and EWS-FLI1 repressed genes by cell line. **E)** Violin plots showing single-cell HOXD13 expression and the IC-EwS EWS-FLI1 signature in the PDX tumors. **F)** Scatter plot of *HOXD13* expression and the IC-EwS EWS-FLI1 signature by PDX. **G)** Scatter plot of HOXD13 activated genes and EWS-FLI1 repressed genes by PDX. Scatter plot of HOXD13 activated genes and EWS-FLI1 repressed genes colored by the mesenchyme development (GO:0060485) gene set score in **H)** EwS cell lines and **I)** PDX tumors. **J)** Summary model of HOXD13 cooperation and antagonism with EWS-FLI1. *r: Pearson’s correlation coefficient*.

Finally, we questioned whether individual cells with differential activity of each of these master transcription factors would harbor differential activation of mesenchymal gene programs. In both cell lines and in PDX tumors, single cells with high HOXD13 activity or high expression of the EWS-FLI1-repressed signature also express high levels of mesenchyme development genes (GO:0060485) (Supplementary Fig. S6G-J). Moreover, integration of the three genesets confirms that individual EwS cells exist along a mesenchymal transcriptional continuum (Fig. 7H, I) and that relative state of individual tumor cells along this axis is determined, at least in part, by the competing activities of HOXD13 and EWS-FLI1 (Fig 7H-J).

## Discussion

Recent studies have shown that successful propagation and metastatic progression of EwS tumors depends on maintaining precise levels of EWS-FLI1 expression and transcriptional activity (57). Moreover, the critical level for tumor growth and progression is dynamic and likely differs at different stages of tumor evolution. For example, while local tumor growth is reliant on continued expression of the fusion, migratory and metastatic properties of EwS cells rely on acquisition of an EWS-FLI1-low state (51, 55, 58, 59). In addition, too much EWS-FLI1 activity is toxic and leads to cell death (57). These observations have led to the premise that EwS cells adhere to the “Goldilocks principle”: they require a dose of oncogene that is “just right” (57). Our current studies have for the first time identified HOXD13 as a key tumor cell-autonomous factor that contributes to maintaining the “just right” level of EWS-FLI1 oncogene activity. In particular, our findings demonstrate that, in addition to cooperating with the fusion at EWS-FLI1-bound and activated loci, HOXD13 serves as an internal rheostat for the EWS-FLI1-repressed signature, creating a state of transcriptional antagonism at target genes that are normally silenced by the fusion. In this manner, EwS cells that harbor high levels of HOXD13 transcriptional activity display key properties of EWS-FLI1-low cells. They express high levels of mesenchymal gene programs, express the mesenchymal stem cell marker CD73 on their cell surface, and have enhanced migratory properties. These results provide a molecular explanation for the observation that HOXD13 loss of function dramatically inhibits EwS metastasis *in vivo* (17). In addition, they underscore the critical importance of tissue-specific transcription factors as key partners for fusion oncoproteins in orchestrating tumor maintenance and progression.

Expression of HOX genes is normally tightly restricted in time and space during embryonic and postnatal life (44, 60). We identified an EWS-FLI1-bound GGAA microsatellite in a highly conserved regulatory region of the posterior HOXD cluster. Targeted epigenomic silencing of this region confirmed that continued expression of *HOXD13* in EwS cells requires EWS-FLI1-mediated binding and activation of this *de novo* GGAA enhancer. However, despite its reproducible capacity for enhancer reprogramming of this site, EWS-FLI1 is insufficient to induce *HOXD13* transcription. In published studies, acute activation of EWS-FLI1 in adult or pediatric human MSCs or human embryonic stem cell-derived neural crest cells failed to induce expression of *HOXD13* (6, 9, 61, 62). However, *HOXD13* mRNA was detected in EWS-FLI1-transduced pediatric MSCs that were cultured in pluripotent stem cell conditions (63) and also in fusion positive human neural crest cells that had been passaged for several weeks (16). Thus, other yet to be defined, cellular and/or microenvironmental cues are needed to complete gene activation following fusion-dependent reprogramming of the HOXD TAD enhancer. It is also noteworthy that the syntenic region in the murine *HoxD* TAD contains a GGAA repeat that is only four repeats in length, well below the “sweet spot” for EWS-ETS binding and activation (64). Consistent with this, we and others have found that EWS-FLI1 does not bind or activate this region or induce *Hoxd13* expression in mouse MSCs (26, 65). We speculate that, given its function as an internal rheostat for EWS-FLI1 activity, HOXD13 may also be required for successful EWS-FLI1-induced malignant transformation. If so, the inability of EWS-ETS proteins to reprogram the murine *Hoxd* TAD may in part explain the continued absence of a genetically engineered mouse model of Ewing sarcoma.

The precise mechanism by which HOXD13 over-rides EWS-FLI1-mediated gene repression remains to be determined but it is unlikely to be exclusively dependent on the absolute levels of each transcription factor or on direct competition between the transcription factors at enhancers. Indeed, although HOXD13 was enriched at GGAA microsatellites and known EWS-FLI1-bound enhancers, these co-bound regions do not directly control mesenchymal genes such as *NT5E* that are repressed in EwS cells. Rather, the two transcription factors appear to compete indirectly to activate (HOXD13) and suppress (EWS-FLI1) mesenchymal gene programs. Given the marked enrichment for wild type ETS, as well as other transcription factor motifs at HOXD13 bound loci, we speculate that the impact of HOXD13 on mesenchymal gene programs, and EWS-FLI1-repressed loci, is mediated through its interactions with other cell context-specific transcription factors beyond the fusion. Wild-type ETS proteins are of particular interest given that they have been previously proposed as modulators of the EWS-FLI1-repressive signature (6). In addition, there is precedence for cooperation between HOX and wild type ETS factors in leukemia and hematopoietic cells where the proteins have been reported to interact and to co-bind at ETS factor motifs (19, 66–68).

In summary, we have discovered that HOXD13 is a direct target of EWS-FLI1 and that co-expression and transcriptional antagonism between these two master transcription factors regulates cell state along a mesenchymal axis. The mechanism of this “competitive cooperation” is mediated directly, by co-binding of the proteins at intronic and intergenic enhancer elements, and indirectly by antagonistic effects on mesenchymal gene programs. In addition, our discovery that EwS cells exist on a transcriptional continuum along a mesenchymal developmental axis may explain why EwS tumors appear to be both histologically and transcriptionally stuck between neuroectodermal and mesodermal cell states (5). Further investigations are now required to fully characterize EwS cell subpopulations and to elucidate if and how tumor cells dynamically shift between among mesenchymal cell states. Ultimately, it will be critical to determine if cell state transitions under the control of master transcription factors such as HOXD13 and EWS-FLI1 contribute to metastatic progression analogous to those conferred by transitional cell states in carcinomas (69).

## Acknowledgements

The authors thank members of the Lawlor lab for helpful discussion, the staff in the University of Michigan Advanced Genomics and Flow cytometry cores for technical assistance, and the staff at the Fred Hutchinson and Seattle Children’s Research Institute cores for flow cytometry, Bioinformatics and Genomics. We thank Phil Corrin and Dolores Covarrubias in Genomics Shared Resource of Fred Hutchinson Cancer Research Center for performing auto CUT&RUN experiments, NGS library preparation and sequencing. We are thankful to Dr. Brian Parkin, who aided in the design and technical completion of the CITE-seq studies.

## Author contributions

AA, JFS, RJHR, and ERL designed the experiments. AA performed wet lab experiments with assistance from JJ, EP, and JP. AA, BM, AGH, SDT, JFS, FW, and SF performed the bioinformatics and statistical analyses for the sequencing data. CH, RJHR, DW, and ML aided in method development and creation and of key reagents. AA and ERL wrote and all authors reviewed and provided input to the final manuscript.

