## Supplementary Methods, Table S1, and Figures for "HOXD13 is a direct EWS-FLI1 target and moderates fusion-dependent transcriptional states"

#### Supplementary Materials

##### Supplementary methods

*Cloning of* UCOE-SFFV-KRAB-dCas9-P2A-mcherry vector

UCOE-SFFV-KRAB-dCas9-P2A vector was created by ligating the AgeI-MluI restriction fragment containing the UCOE and SFFV promoter from UCOE-SFFV-dCas9-KRAB-BFP (Addgene, 85969) into the complementary fragment of pHR-SFFV-KRAB-dCas9-P2A-mCherry (Addgene, 60954)(52). UCOE-SFFV-KRAB-dCas9-P2A-mcherry lentivirus was transduced into cells in serum free media + 5ug/uL polybrene for 6 hours followed by addition of complete media (1:1).

*gRNA design and cloning*

The sgRNAs were designed with Bsmb1 cut sites, cloned into the sgOpti vector (Addgene, 85681), and validated by sanger sequencing. The negative control sgRNA NC_chr1 targets a unique intergenic site of no known function on chromosome 1, and showed no significant functional effect in a prior CRISPRi screen.

Cells transduced with UCOE-SFFV-KRAB-dCas9-P2A-mcherry were FACS sorted twice on mCherry expression. sgRNAs (Supplementary Table S1) were designed against nucleosome free regions flanking GGAA repeat sites using the Broad institutes GPP sgRNA designer. sgRNAs with high on-target scores were chosen and cloned into the sgOpti vector (Addgene, 85681). Lentiviral sgRNA-containing sgOpti vectors were transduced into stably expressing dCas9-KRAB-mcherry cells and puromycin selected before collection after 8 days.

**Immunofluorescence**

Frozen sections of E13.5 *Hoxd13* WT, heterozygous, and knockout embryos (26) were formalin fixed, frozen in OCT and sectioned. Slides were thawed at RT, washed with PBS, blocked for 1 hour at room temperature (10% donkey serum + 0.2% triton) and incubated with the HOXD13 primary antibody in blocking buffer overnight at 4C. Slides washed with 0.2% triton PBS before addition of donkey anti-rabbit 488 secondary in blocking buffer for 1 hour at RT. After three more PBS + 0.2% triton washes 1:1000 DAPI (Fisher) was added. Images were taken using an inverted Olympus IX83 (Tokyo, Japan). CellSens Dimensions captured the images. For immunocytochemistry, cells were fixed in 4% paraformaldehyde for 15 minutes and permeabilized in 0.5% Triton in PBS for 10 minutes. After washes in PBS, cells were blocked for 1 hour at RT in 0.2% BSA + 5% Goat Serum and then incubated either HOXD13 or IgG or 1 hour at RT in blocking buffer. Following 3 washes, a fluorescent secondary antibody was added and incubated for 1 hour at RT, followed by washes and DAPI (1:10 000) incubation. Images were taken on a Lecia DMi8 microscope using the Lecia software.

**Mapping mouse enhancers to human**

To map mouse enhancer sites in the regulatory HOXD domain the UCSC LIFTOVER tool was used to go from mm9 to hg19. Five developmental enhancer locations mapped (hg19): Island I (chr2:176124212-176134930), Island II (chr2:176237060-176246196), Island III (chr2:176423732-176424979), Island IV (chr2:176503826-176503873), Island V (chr2:176549574-176555217). PHE GGAA coordinates (hg19): chr2:176268550-176268900.

**Bru-seq/RNA-seq and analysis**

Cell line samples were prepared as described previously (28). Bru-seq was performed on A673 and CHLA10 cells following HOXD13 knockdown. In brief, 72 hours after doxycycline addition to either shNS or shHOXD13#1 cells bromouridine (2mM; Sigma) was added for 30 minutes. Nascent RNA was immunoprecipitated out and cDNA libraries were prepared and sequenced using an Illumina HiSeq 2500 for 50-bp single-end reads at the University of Michigan Sequencing Core. Reads were processed, mapped (hg38), and quantified using the Bru-seq analysis pipeline (28). Poly(A)-capture RNA-seq was performed for TC32 HOXD13 knockdown cells. Libraries were prepared with NEBNext Ultra II RNA Library Prep kit and Paired end 150 bp sequencing was performed on a Novaseq600 by Novogene. Adapter Trimming was performed using Trim Galore (Babraham Institute). Trimmer reads were aligned to GRCh38 using the STAR aligner (29). Differential expression was calculated using DESeq2 v1.18.1 (30) using an FDR-adjusted *p*-values <0.05. Overrepresentation analysis was performed using a Fisher's exact test to quantify the overlap of differentially expressed genes with the gene sets in the Broad Institute's Molecular Signatures Database (MSigDB) (31). Heatmaps and volcano plots were made with the R packages pheatmap and EnhancedVolcano, respectively.

**Automated CUT&RUN Sequencing and analysis**

250-500K cells/reaction were collected and washed with Wash Buffer (20 mM HEPES-NaOH pH 7.5, 150 mM NaCl, 0.5 mM Spermidine, Protease Inhibitor) and incubated for 5 min at room temperature with BioMag Plus Concanavalin A-coated magnetic beads

(Bangs Laboratories, BP531; 15 uL/ reaction). Supernatant was removed and beads/cells were resuspended in antibody buffer (20 mM HEPES-NaOH pH 7.5, 150 mM NaCl, 0.5 mM Spermidine, 0.05% Digitonin, 2 mM EDTA, Protease Inhibitor) and antibodies were added (Supplementary Table S1). Samples were incubated 24-48 hours at 4°C then submitted to the Genomics core at the Fred Hutchinson and the Automated CUT&RUN protocol was performed as described (32)(https://www.protocols.io/view/autocut-run-genome-wide-profiling-of-chromatin-pro-ufeetje).

FastQC 0.1.9 (https://www.bioinformatics.babraham.ac.uk/projects/fastqc/) was used to examine read quality and paired-end reads were aligned to hg38 using Bowtie2.3.5. (33). Peaks of histone marks were called using MACS2.2.7 (34) with both treatment-only and IgG-controlled modes. Narrow peaks were called and filtered for H3K4me1, H3K4me3, and H3K27ac, while broad peaks were called and filtered for H3K27me3. For downstream analyses, IgG-controlled narrow peaks were kept if they overlapped with treatment-only peaks from the same sample, q-value is less than 0.01 and fold enrichment is over 2; treatment-only broad peaks were kept if they overlap with IgG-controlled peaks from the same sample, peak width is at least 2 Kbp, q-value is less than 0.01 and fold enrichment is over 2.

For HOXD13, in addition to MACS2 narrow peak calling and peak filtering described above, SEACR (35) was also run with stringent mode. Filtered MACS2 peaks that overlap with SEACR peaks were passed on to the downstream analyses. BEDTools 2.30.0 (36) was used to identify overlapping peaks between marks. Peak annotation and motif analysis was performed with HOMER 4.11 (37). HOMER was also used to identify TF motifs in peaks. Overlaps of genes were calculated using the Fischer's exact test from the GeneOverlap package and overlaps of genomic sites were calculated using ChIPpeakAnno. Length and number of GGAA/CCTT sites were calculated for overlapped HOXD13 and EWS-FLI1 binding sites within 250bp up and downstream the peak in hg38.

**CITE-seq processing and analysis**

CITE-seq allows for matched transcript and cell surface antigen profiling of individual cells (38, 39). 500 000 CHLA10 and A673 cells were resuspended in PBS + 1% FBS and spun at 300g for 10 minutes at 4C and then resuspended in 25 uL Biolegend staining buffer (420201, San Diego, CA). 2.5 uL human TruStain FcX (422301, San Diego, CA) was added per sample and incubated at 4C for 10 minutes. Hash-Tag Antibodies, used to label the samples and minimize batch effects, were added at 250 ng/sample (0.5 uL/sample) and incubated at 4C for 30 minutes. Cells were washed twice in 1 mL staining buffer and spun at 300g for 5 minutes at 4C. Samples were pooled 1:1 at 1 000 cells/uL and libraries were generated using the 3' V3 10X Genomics Chromium Controller following the manufacturer's protocol (CG000183, Pleasanton, CA). Final library quality was assessed using the Tapestation 4200 (Agilent, Santa Clara, CA) and libraries were quantified by Kapa qPCR (Roche). Pooled libraries were then subjected to paired-end sequencing according to the manufacturer's protocol (Illumina NovaSeq 6000). Bcl2fastq2 Conversion Software (Illumina) was used to generate de-multiplexed Fastq files and the CellRanger (3.1) Pipeline (10X Genomics) was used to align reads and generate count matrices. Analysis performed using Seurat (40) and Monocle 3 (41). A graph-based unsupervised clustering approach (40) followed by the nonlinear dimensionality reduction technique, Uniform Manifold Approximation Projection (UMAP) was then used to cluster and visualize the data within each cell line (42).

**Analysis of EWS-FLI1 ChIP-seq for hg38 alignment**

Raw fastqs from GEO were downloaded using SRA tools (GSE61953)(6). These were converted to BED files using chromap aligned to hg38 and then into bigwigs using Kent Utilities from the UCSC tools package.

**Statistical analysis**

Unless otherwise indicated, data were analyzed using GraphPad Prism software version 9.0 (GraphPad Software, San Diego, CA, USA). All statistical analyses were performed with Student's t-test / One-way ANOVA followed by Tukey multiple comparison test / or Two-way ANOVA followed by Sidak's multiple comparison test. Data are expressed as means and SEM from at least three independent experiments. Asterisk denoting $p<0.05$ (*) or $p<0.01$ (**).

#### Supplementary information
#### Table S1 Supplementary materials

| REAGENT or RESOURCE | SOURCE | IDENTIFIER | Concentration |
|---|---|---|---|
| **Antibodies** | | | |
| Rabbit polyclonal anti-FLI1 | abcam | Cat# ab15289; RRID:AB_301825 | ChIP- 5ug/1x10^6 cells; WB- 5ug/2.5 mL in 5% milk & TBST |
| Rabbit polyclonal anti-HOXD13 | This study | N/A | WB- 1:500 in 5% milk & TBST; ICC- 1:100 (4.6 ug) 10% donkey serum + 0.2% triton & PBS; CUT&RUN: 1:20 |
| Rabbit monoclonal IgG (ICC) | Abcam | Cat# ab172730; RRID:AB_2687931 | ICC- 4.6 ug "same as above"" |
| Rabbit monoclonal ant-GAPDH | Cell signaling | Cat# 2118; RRID:AB_561053 | WB- 1:1 000 in 5% milk & TBST |
| Mouse monoclonal anti-GAPDH | Invitrogen | Cat# AM4300; RRID:AB_2536381 | WB- 1:13 000 in 5% milk & TBST |
| Rabbit polyclonal anti-histone H3K27ac (ChIP-RTqPCR) | abcam | Cat# ab4729; **RRID:AB_2118291** | ChIP- 2ug/1x10^6 cells |
| Rabbit monoclonal anti-histone H3K27ac (CUT&RUN) | Millipore | Cat# MABE647; RRID:AB_2893037 | CnR- 1:50 |
| Rabbit polyclonal anti-histone H3K4me1 (ChIP-RTqPCR) | abcam | Cat# ab8895; RRID:AB_306847 | ChIP- 2ug/1x10^6 cells |
| Rabbit polyclonal anti-histone H3K4me1 (CUT&RUN) | Epicypher | Cat# 13-0040 | CnR- 1:100 |
| Rabbit polyclonal anti-histone H3K9me3 (ChIP-RTqPCR) | abcam | Cat# ab8898; RRID:AB_306848 | ChIP- 2ug/1x10^6 cells; CnR- 1:100 |
| Rabbit polyclonal anti-histone H3K4me3 (CUT&RUN) | Active motif | Cat# 39159; RRID:AB_2615077 | CnR- 1:50 |
| Rabbit monoclonal anti-histone H3K27me3 (CUT&RUN) | Cell Signaling | Cat# 9733; RRID:AB_2616029 | CnR- 1:100 |
| Rabbit polyclonal IgG (ChIP-RTqPCR,CUT&RUN) | abcam | Cat# ab37415; RRID:AB_2631996 | ChIP- 2-5ug/1 million cells<br>CnR- 1:50 |
| Mouse monoclonal anti-CD73 BB515 | BD Pharmingen | Cat# BDB565110; RRID:AB_2739072 | Flow- 5 uL/1x10^6 cells |
| Mouse monoclonal anti-CD271 BB515 | BD Pharmingen | Cat# BDB564580; RRID:AB_2738853 | Flow- 5 uL/1x10^6 cells |
| Mouse IgG1, kappa, BB515 | BD Pharmingen | Cat# BDB564416; RRID:AB_2721017 | Flow- 5 uL/1x10^6 cells |
| Mouse monoclonal anti-CD73 PE | BD Pharmingen | Cat# BDB561014; RRID:AB_2033967 | Flow- 20 uL/1x10^6 cells |
| Mouse monoclonal anti-CD271 PE | BD Pharmingen | Cat# BDB560927; RRID:AB_10564069 | Flow- 20 uL/1x10^6 cells |
| Mouse IgG1, kappa, PE | BD Pharmingen | Cat# BDB555749; RRID:AB_396091 | Flow- 20 uL/1x10^6 cells |
| TotalSeq-A0251 anti-human hashtag 1 (HTO1) | BioLegend | Cat# 394601; RRID:AB_2750015 | CITE- 1 uL/1x10^6 cells |
| TotalSeq-A0252 anti-human hashtag 2 (HTO2) | BioLegend | Cat# 394603; RRID:AB_2750016 | CITE- 1 uL/1x10^6 cells |
| IRDye 800CW Goat anti-Rabbit IgG Secondary Antibody | LiCor | Cat# 926-32211 | WB- 1:10, 00 in TBST |
| IRDye 680CW Goat anti-Rabbit IgG Secondary Antibody | LiCor | Cat# 926-68071 | WB- 1:10 000 in TBST |
| IRDye 680CW Goat anti-Mouse IgG Secondary Antibody | LiCor | Cat# 926-68070 | WB- 1:10 000 in TBST |
| Donkey anti-Rabbit 488 Secondary | Fisher | Cat# A11034 | IF- 1:500 |

| | | | |
|---|---|---|---|
| Alexa Fluor 647 Goat anti rabbit IgG | Life Technologies | Cat#:A-21245 | ICC- 1:1 000 |
| **Deposited data** | | | |
| Bru-seq, RNA-seq, CUT&RUN, and CITE-seq | This study | GSE182513 | N/A |
| CITE-seq analysis | This study | https://github.com/LawlorLab/HOXD13-Paper | N/A |
| EWS-FLI1 knockdown bulk RNA-seq | Riggi et al., 2014 | GSE61953 | N/A |
| EWS-FLI1, H3K4me1, H3K4me3, H3K327me3 ChIP-seq in A673 and SKNMC cells and primary tumors. | Riggi et al., 2014 | GSE61953 | N/A |
| ATAC-seq and H3K27ac and H3K4me1 ChIP-seq in MSCs | Riggi et al., 2014 | GSE61953 | N/A |
| EWS-FLI1, H3K4me1 ChIP-seq data and ATAC-seq in MSCs | Boulay et al., 2017 | GSE94275 | N/A |
| EWS-FLI1, H3K4me1, H3K4me3, H3K27me3 ChIP-seq data in MSCs | Boulay et al., 2018 | GSE106925 | N/A |
| H3K27ac and H3K4me1 in Osteosarcoma primary tumors | Morrow et al., 2018 | GSE74230 | N/A |
| Single-cell RNA sequencing from A673 shEF xenografts and PDX tumors | Aynaud et al., 2020 | GSE130025 | N/A |
| **Oligonucleotides** | | | **Size (bp) & location hg19** |
| Taqman probe human 18S (housekeeping) | Thermo | Hs03003631_g1 | 69 bp |
| Taqman probe human B2M | Thermo | Hs00984230_m1 | 81 bp |
| Taqman probe human HOXD13 | Thermo | Hs00968515_m1 | 102 bp |
| Taqman probe human HOXD11 | Thermo | Hs00360798_m1 | 133 bp |
| Taqman probe human HOXD10 | Thermo | Hs00157974_m1 | 61 bp |
| Taqman probe EWS-FLI1 type 1 fusion | Thermo | Hs03024497_ft | 80 bp |
| Taqman probe NR0B1 | Thermo | Hs00230864_m1 | 92 bp |
| Taqman probe VRK1 | Thermo | Hs00177470_m1 | 85 bp |
| VRK1 GGAA enhancer- *EWS-FLI1 binding* F: (5'ACTCGGCTTTTTGCAACTTC 3') | Riggi et al. 2014 | N/A | 135; chr14:97681757-97681891 |
| VRK1 GGAA enhancer- *EWS-FLI1 binding* R: (5' CCTCTTGCCTTCCTTCCTTC 3') | Riggi et al. 2014 | N/A | |
| VRK1 GGAA enhancer- *histone marks* F: (5' CGATGGGTGATCAATGAGTG 3') | This study | N/A | 236; chr14:97681017-97681252 |
| VRK1 GGAA enhancer- *histone marks* R: (5' AAGAGAGCTTGGGGAGGAAG 3') | This study | N/A | |
| Negative control region in chr2 F: (5' ATGGTGATTCTCAGCCTCCA 3') | This study | N/A | 107; chr2:176644216-17664432 |
| Negative control region in chr2 R: (5' TGCAGGATTTAAGGGAACCA 3") | This study | N/A | |
| PHE- *EWS-FLI1 binding* F: (5' CTCCCTTGCTTCCTTCCTTC 3') | This study | N/A | 104; chr2:176268617-176268720 |
| PHE- *EWS-FLI1 binding* R: (5' TACCCAGAAACTGGCACACA 3') | This study | N/A | |
| PHE- *histone marks* F: (5' AAATGGCATGATCACTTTTGTG 3') | This study | N/A | 92; chr2:176269105-176269196 |
| PHE- *histone marks* R: (5' CTTTCTCTGCCCCCAGTTTT 3') | This study | N/A | |
| SOX2 GGAA enhancer F: (5' GAAGTGCACCCTATGCCAGT 3') | Riggi et al. 2014 | N/A | 104; chr3:181901578-181901681 |
| SOX2 GGAA enhancer | Riggi et al. 2014 | N/A | |

| | | | |
|---|---|---|---|
| R: (5' TCCTCTGTGGGGGTTATCCA 3') | | | |
| **sgRNAs for Crispri** | | | |
| Negative control-chr1 non-targeting site sgRNA<br>F: (5' CACCGCCAAACATTCCAGCTATCCA 3')<br>*red indicates restriction cut sites | Fulco et al. 2016 | N/A | |
| Negative control-chr1 non-targeting site sgRNA<br>R: (5' AAACTGGATAGCTGGAATGTTTGGC 3')<br>*red indicates restriction cut sites | Fulco et al. 2016 | N/A | |
| SOX2 GGAA sgRNA<br>F: (5' CACCTTATCCATCTAACAGGTGGG 3') | Boulay et al. 2018 | N/A | |
| SOX2 GGAA sgRNA<br>R: (5' AAACCCCACCTGTTAGATGGATAA 3') | Boulay et al. 2018 | N/A | |
| PHE_1 sgRNA<br>F: (5' CACCGTGTCCCTAAGTATACAGATA 3') | This study | N/A | |
| PHE_1 sgRNA<br>R: (5' AACTATCTGTATACTTAGGGACAC 3') | This study | N/A | |
| PHE_2 shRNA<br>F: (5' CACCGGCCCTACTATCTCTCAGTGA3') | This study | N/A | |
| PHE_2 sgRNA<br>R: (5' AACTCACTGAGAGATAGTAGGGCC 3') | This study | N/A | |
| PHE_3 sgRNA<br>F: (5' CACCGATGTCATGTATAATCCTGCA 3') | This study | N/A | |
| PHE_3 sgRNA<br>R: (5' AACCTAGATTACAAATTCCGTGCC 3') | This study | N/A | |
| **Recombinant DNA** | | | |
| pCD/NL-BH*DDD | addgene | Cat# 17531 | N/A |
| pMD2.G | addgene | Cat# 12259 | N/A |
| pLKO.1 non-targeting shRNA | Sigma | Cat# SHC002 | N/A |
| pLKO.1 FLI1-targeting shRNA | Sigma | TRCN0000005322 | N/A |
| pTripz shNS | Horizon-Dharmacon | RHS4743 | N/A |
| pTripz shHOXD13 #1 | Horizon-Dharmacon | RHS4696-200755961, clone: V3THS_321416 | N/A |
| pTripz shHOXD13 #2 | Horizon-Dharmacon | RHS4696-200680827, clone: V2THS_93475). | N/A |
| pCLS-EGFP empty | | | N/A |
| EWS-FLI1-V5-2A-EGFP | | | N/A |
| pLIV empty | M. Rivera Laboratory, MGH, Boston, MA, USA | Riggi et al., 2014 | N/A |
| pLIV EWS-FLI1 | M. Rivera Laboratory, MGH, Boston, MA, USA | Riggi et al., 2014 | N/A |
| UCOE-SFFV-dCas9-KRAB-BFP | Addgene | 85969 | N/A |
| pHR-SFFV-KRAB-dCas9-P2A-mCherry | Addgene | 60954 | N/A |
| UCOE-SFFV-KRAB-dCas9-P2A | This study | N/A | N/A |
| sgOpti | Addgene | 85681 | N/A |

### Supplementary Figures

#### Figure S1

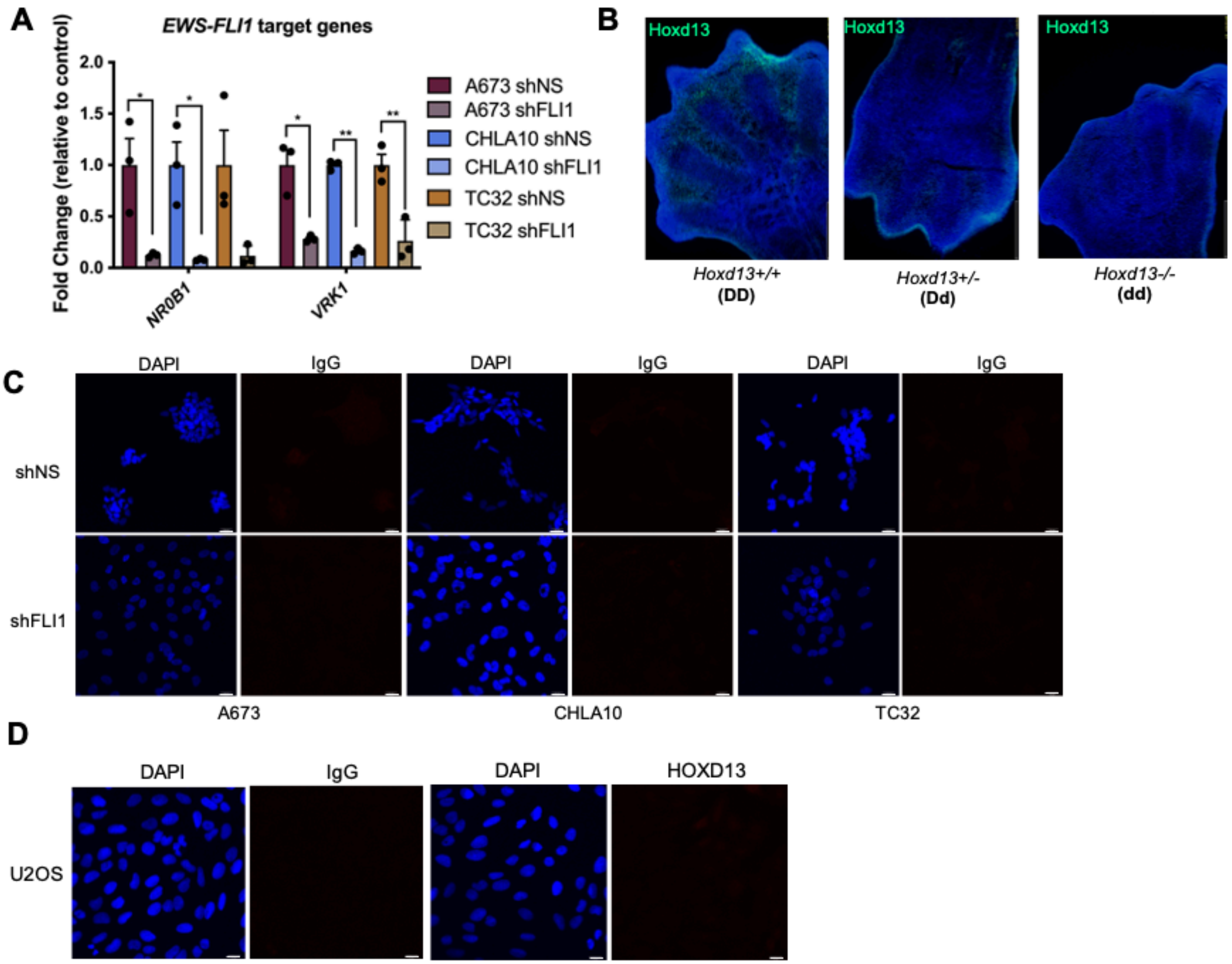

**Figure S1. Authentication of EWS-FLI1 knockdown cells and the HOXD13 antibody**

A) qRT-PCR of EWS-FLI1 target genes, *NR0B1* and *VRK1* after knockdown. Error bars representative of SEM from three independent replicates. * $p<0.05$; ** $p<0.01$; Two-way ANOVA; Sidak's multiple comparison test.

B) Immunofluorescence detection of Hoxd13 protein expression in distal limbs (autopods) derived from E13.5 *Hoxd13* WT (+/+), het (+/-), or knockout (-/-) mouse embryos.

C-D) Immunofluorescent staining of shFLI1 and U2OS cells with IgG isotype control and HOXD13 antibodies, respectively (secondary antibody Alexa647). Nuclear counterstain was performed with DAPI. Scale bar is 10 um.

### Figure S2

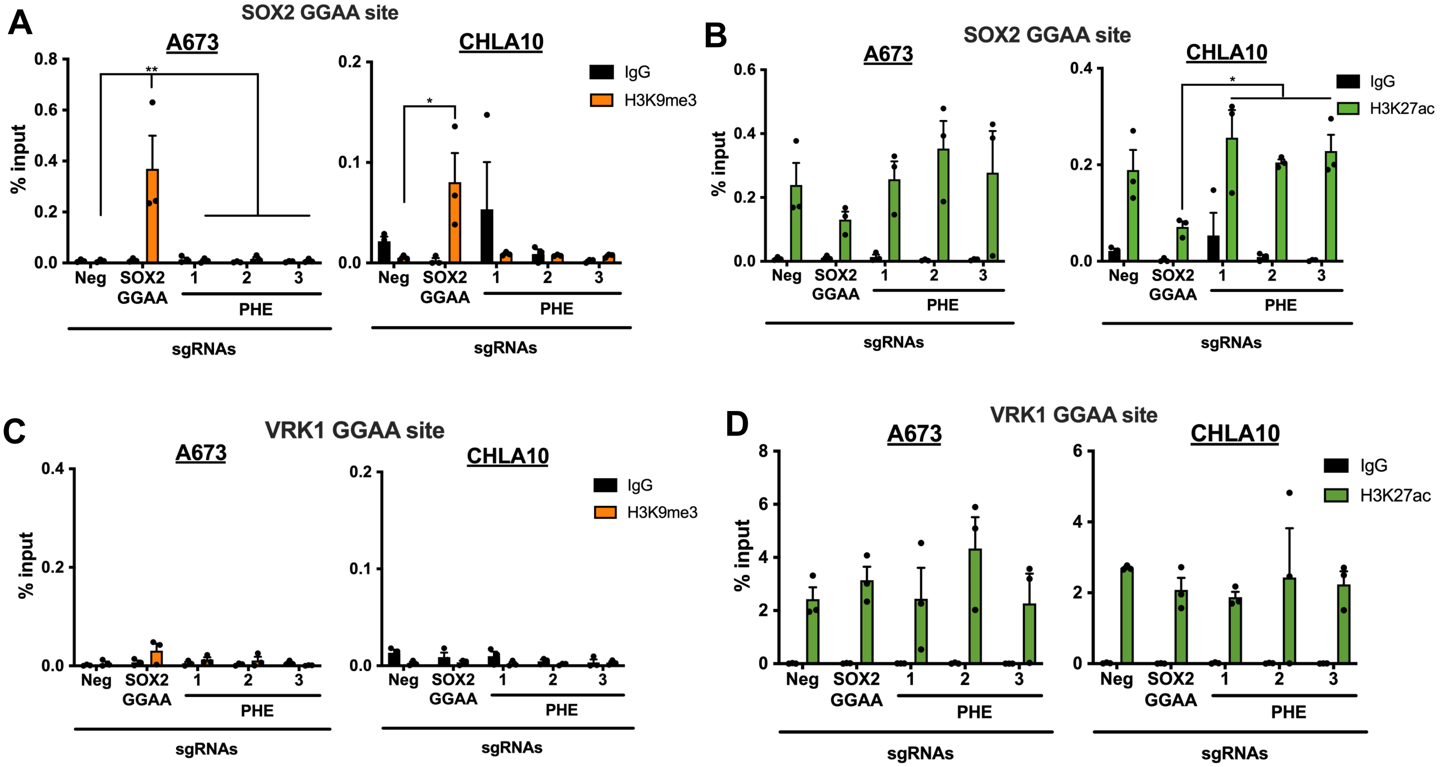

A
SOX2 GGAA site
A673
CHLA10
IgG
H3K9me3
% input
Neg
SOX2
GGAA
1
2
3
PHE
sgRNAs
B
SOX2 GGAA site
A673
CHLA10
IgG
H3K27ac
C
VRK1 GGAA site
A673
CHLA10
IgG
H3K9me3
D
VRK1 GGAA site
A673
CHLA10
IgG
H3K27ac

**Figure S2. Enhancer RNA expression at the PHE**

ChIP-qPCR for A) H3K9me3 and B) H3K27ac at the SOX2 GGAA enhancer site. ChIP-qPCR for C) H3K9me3 and D) H3K27ac at the VRK1 GGAA enhancer site. VRK1 enhancer used as a control GGAA site. Error bars are representative of SEM from at least three independent experiments. * $p<0.05$; ** $p<0.01$; Two tailed t-test; Two-way ANOVA; Sidak's multiple comparison test.

### Figure S3

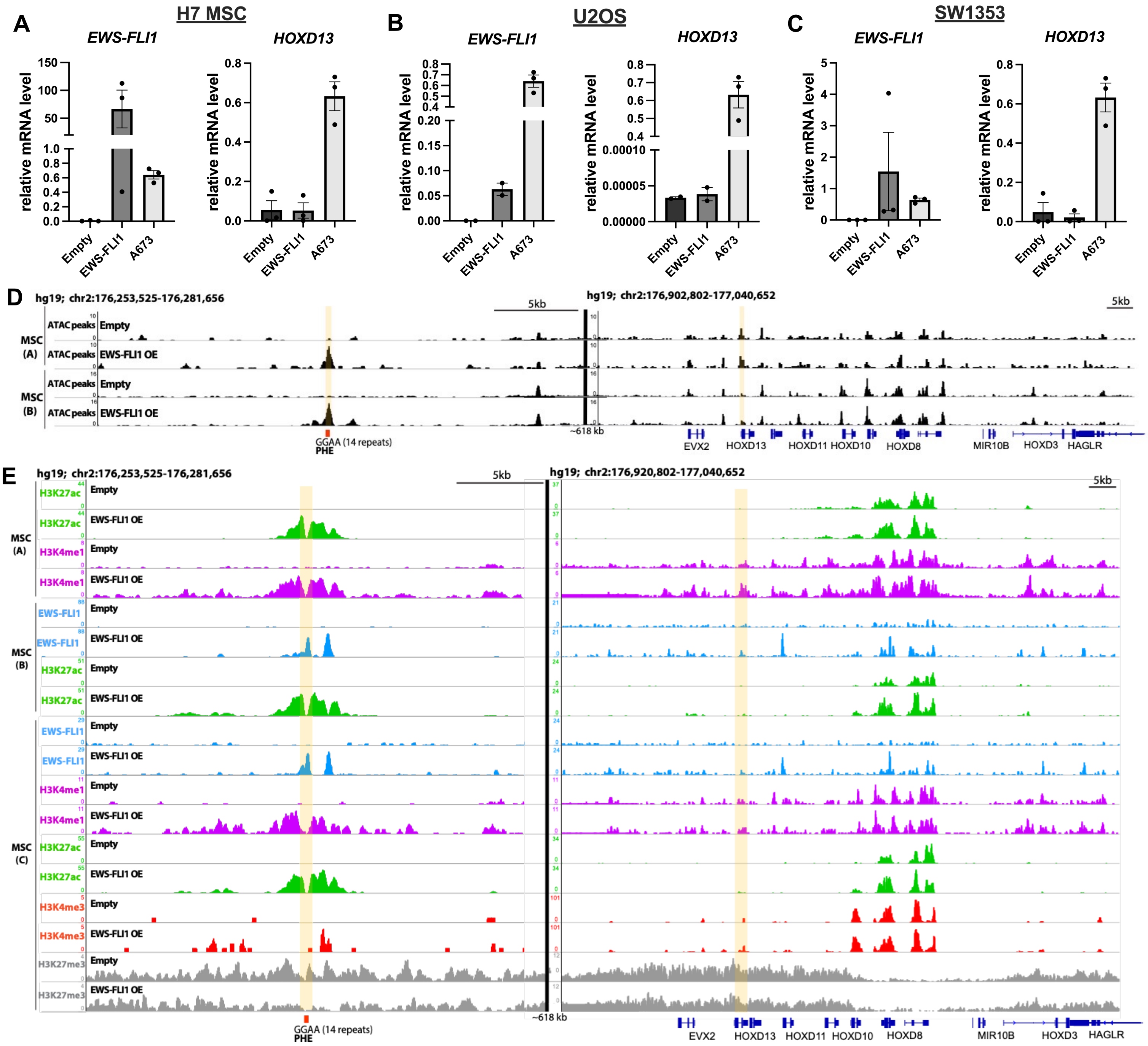
A
H7 MSC
EWS-FLI1
HOXD13
relative mRNA level
Empty
EWS-FLI1
A673
B
U2OS
C
SW1353
D
hg19; chr2:176,253,525-176,281,656
hg19; chr2:176,902,802-177,040,652
5kb
MSC (A)
MSC (B)
ATAC peaks
Empty
EWS-FLI1 OE
GGAA (14 repeats)
PHE
~618 kb
EVX2
HOXD13
HOXD11
HOXD10
HOXD8
MIR10B
HOXD3
HAGLR
E
hg19; chr2:176,253,525-176,281,656
hg19; chr2:176,920,802-177,040,652
MSC (C)
H3K27ac
H3K4me1
H3K4me3
H3K27me3

**Figure S3. EWS-FLI1 is not sufficient to induce *HOXD13* transcription in heterologous cell types**

qRT-PCR measurement of *EWS-FLI1* and *HOXD13* expression in A) H7 MSCs, B) U20S, and C) SW1353 cells after EWS-FLI1 transduction. A673 cells shown for comparison.

D) ATAC-seq tracks at the PHE and HOXD locus in MSCs transduced with EWS-FLI1 or an empty vector.

E) ChIP-seq tracks for EWS-FLI1, H3K27ac, H3K4me1, H3K4me3, and H3K27me3 at the PHE and HOXD locus in MSC's overexpressing EWS-FLI1 or an empty vector. MSC ATAC and ChIP-seq tracks from Refs. (A-C: (6), (9), (10), respectively).

### Figure S4

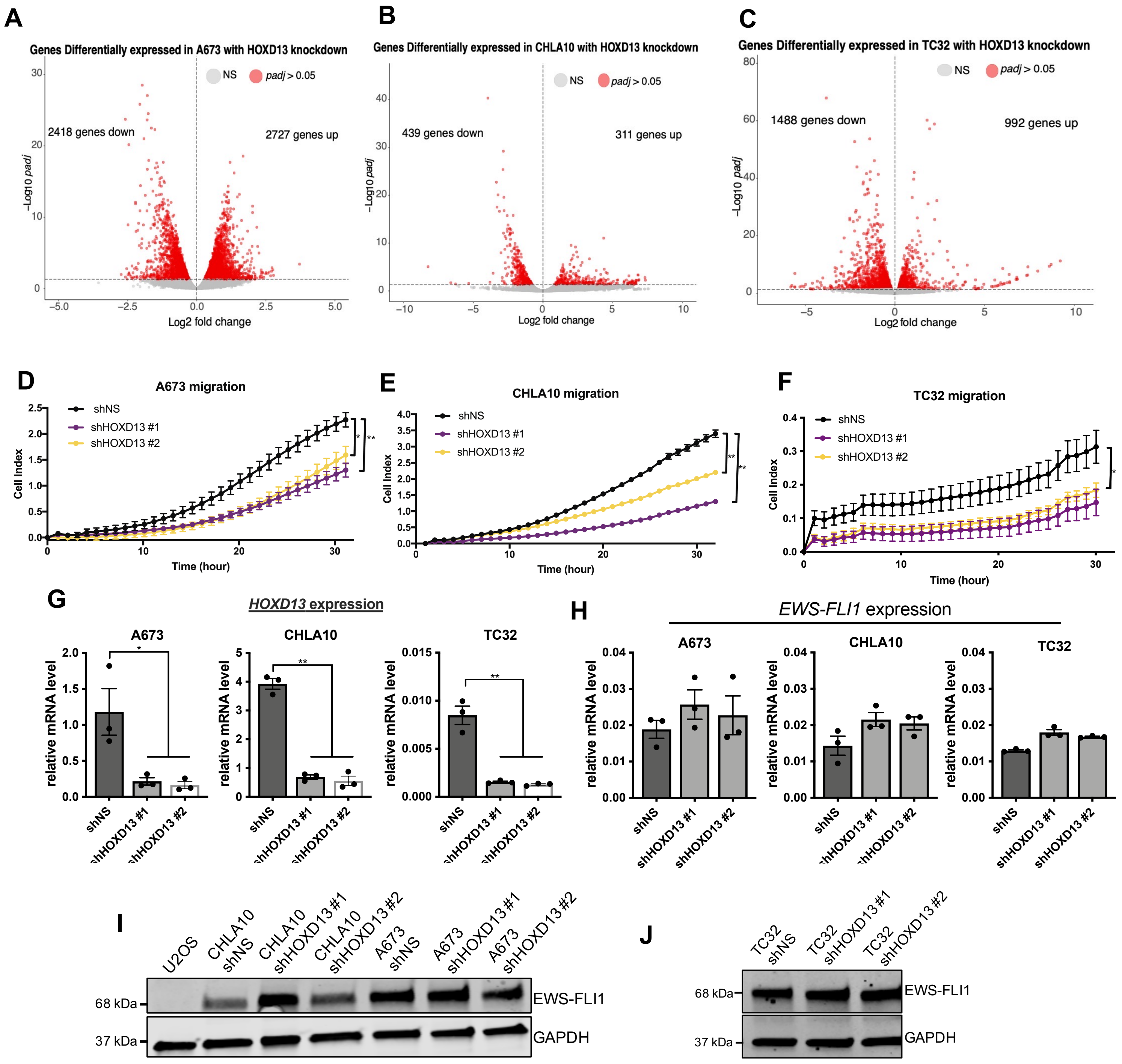
A
Genes Differentially expressed in A673 with HOXD13 knockdown
NS
padj > 0.05
2418 genes down
2727 genes up
−Log10 padj
Log2 fold change
B
Genes Differentially expressed in CHLA10 with HOXD13 knockdown
439 genes down
311 genes up
C
Genes Differentially expressed in TC32 with HOXD13 knockdown
1488 genes down
992 genes up
D
A673 migration
shNS
shHOXD13 #1
shHOXD13 #2
Cell Index
Time (hour)
E
CHLA10 migration
F
TC32 migration
G
HOXD13 expression
A673
CHLA10
TC32
relative mRNA level
H
EWS-FLI1 expression
I
U2OS
CHLA10 shNS
CHLA10 shHOXD13 #1
CHLA10 shHOXD13 #2
A673 shNS
A673 shHOXD13 #1
A673 shHOXD13 #2
68 kDa
37 kDa
EWS-FLI1
GAPDH
J
TC32 shNS
TC32 shHOXD13 #1
TC32 shHOXD13 #2

**Figure S4. HOXD13-regulated gene programs are largely cell line-dependent**

A-C) Volcano plots for each cell line depicting the differentially expressed genes after HOXD13 knockdown (*padj <0.05*).

D-F) xCELLigence Real Time Cell Migration assay of control and HOXD13 knockdown cells. Error bars are representative of SEM from at least two independent experiments with 5 technical replicates per condition. * $p<0.05$; ** $p<0.01$

G) qRT-PCR of *HOXD13* expression of control (shNS) and *HOXD13* (shHOXD13) knockdown cells.

H) qRT-PCR of EWS-FLI1 expression in HOXD13 knockdown cells.

I-J) Western blot of EWS-FLI1 expression in HOXD13 knockdown cells. Two-tailed *t*-test; One-way ANOVA followed by a Tukey's multiple comparison test. Error bars representative of SEM from three independent replicates. * $p<0.05$; ** $p<0.01$; Two-way ANOVA; Sidak's multiple comparison test.

### Figure S5

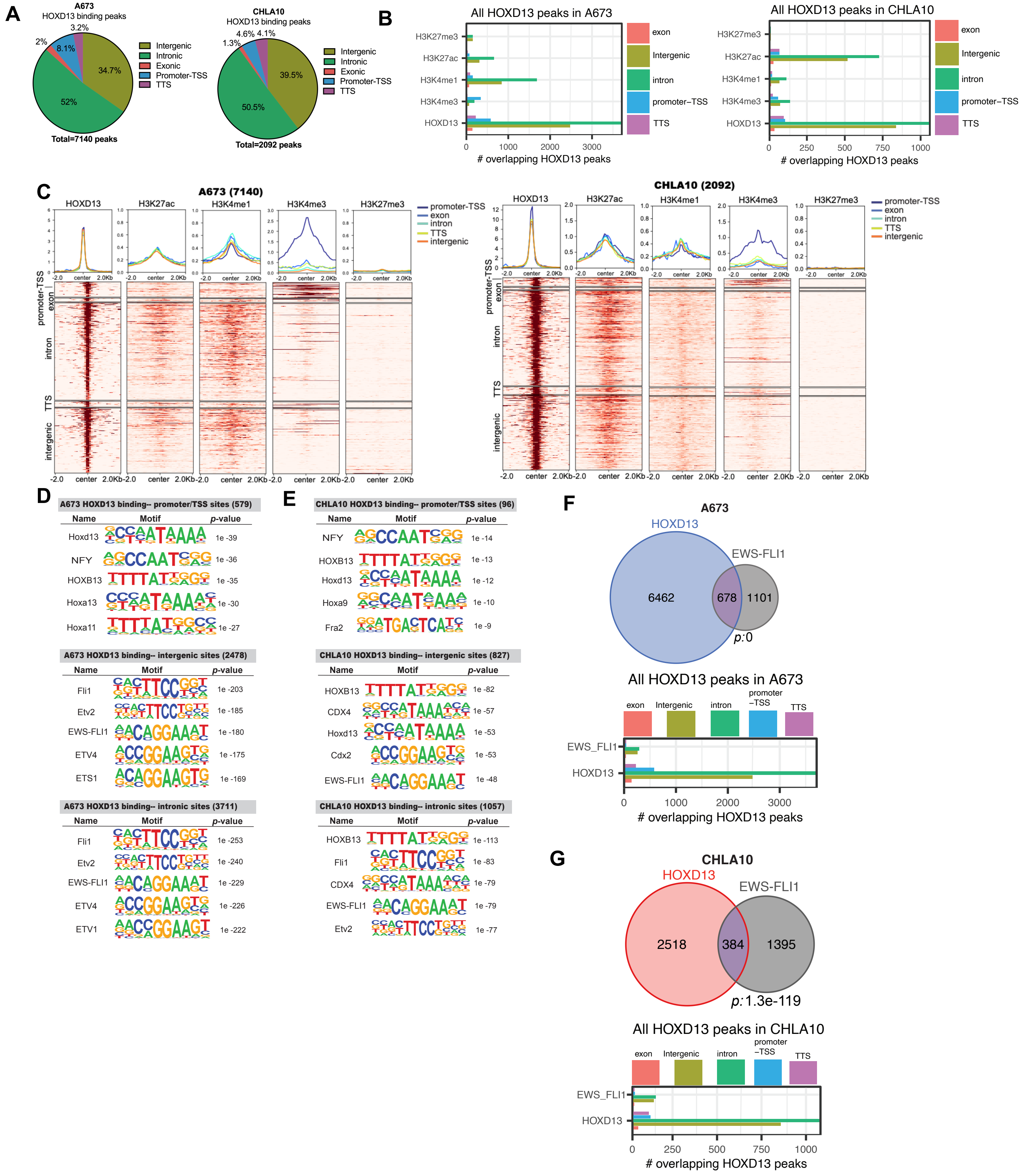

A
A673
HOXD13 binding peaks
Intergenic
Intronic
Exonic
Promoter-TSS
TTS
Total=7140 peaks
CHLA10
HOXD13 binding peaks
Total=2092 peaks
B
All HOXD13 peaks in A673
All HOXD13 peaks in CHLA10
### overlapping HOXD13 peaks
C
A673 (7140)
CHLA10 (2092)
HOXD13
H3K27ac
H3K4me1
H3K4me3
H3K27me3
promoter-TSS
exon
intron
TTS
intergenic
D
A673 HOXD13 binding-- promoter/TSS sites (579)
A673 HOXD13 binding-- intergenic sites (2478)
A673 HOXD13 binding-- intronic sites (3711)
E
CHLA10 HOXD13 binding-- promoter/TSS sites (96)
CHLA10 HOXD13 binding-- intergenic sites (827)
CHLA10 HOXD13 binding-- intronic sites (1057)
F
A673
HOXD13
EWS-FLI1
6462
678
1101
p:0
All HOXD13 peaks in A673
G
CHLA10
HOXD13
EWS-FLI1
2518
384
1395
p:1.3e-119
All HOXD13 peaks in CHLA10

**Figure S5. HOXD13 binding distribution is similar in A673 and CHLA10 cells**

A) Pie chart showing the genomic distribution HOXD13 binding sites in A673 and CHLA10 cells.

B) Bar chart summarizing shared binding sites and associated histone marks at these sites.

C) Tornado plots depicting shared HOXD13 binding by genomic location and the associated histone marks at these sites.

HOMER Motif analysis by genomic location for the HOXD13 binding sites in D) A673 and E) CHLA10 cells.

Venn diagrams showing the overlap between HOXD13 binding sites and published EWS-FLI1 binding sites in F) A673 and G) CHLA10.

### Figure S6

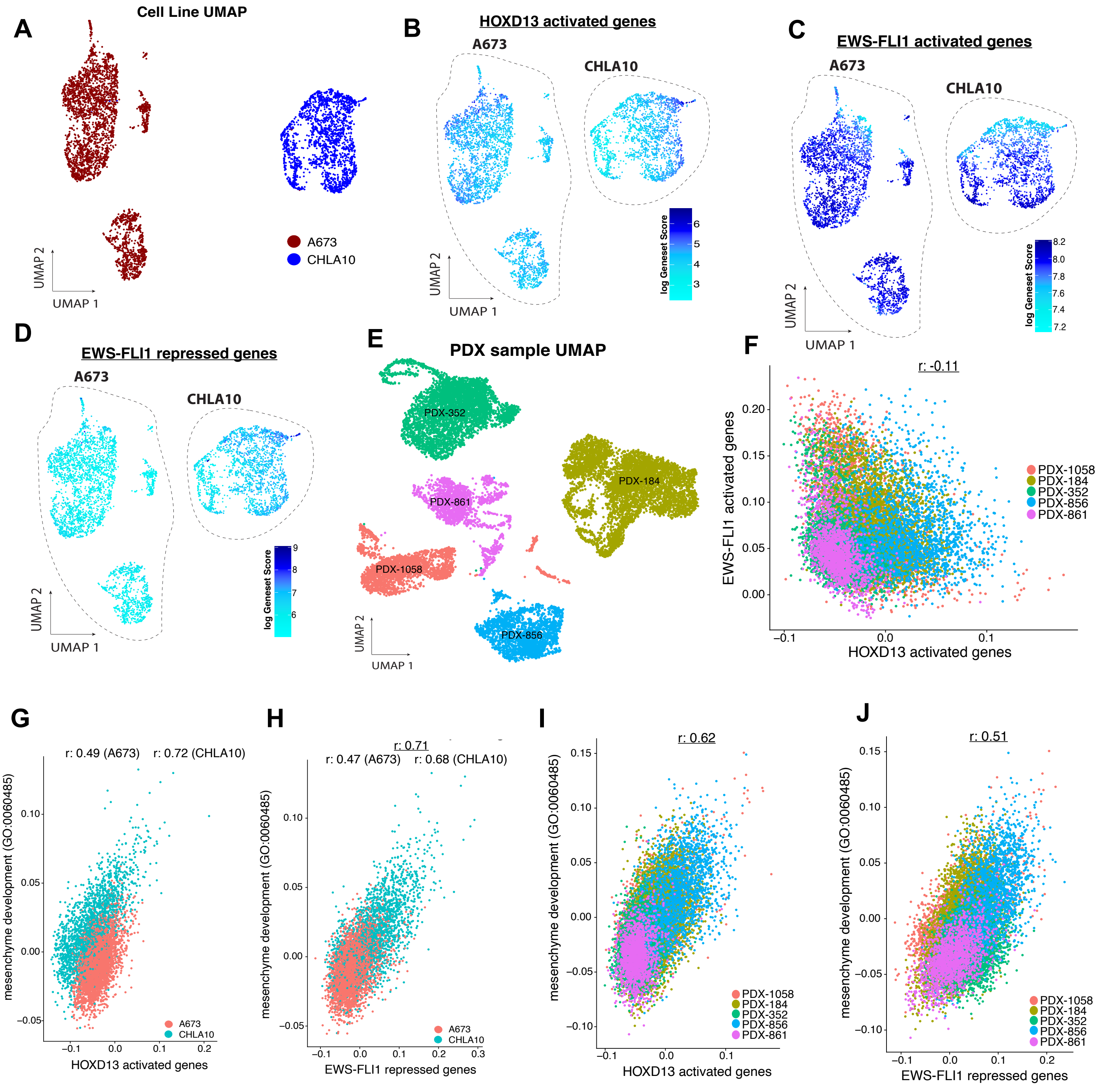

A
Cell Line UMAP
A673
CHLA10
UMAP 1
UMAP 2
B
HOXD13 activated genes
A673
CHLA10
log Geneset Score
C
EWS-FLI1 activated genes
A673
CHLA10
log Geneset Score
D
EWS-FLI1 repressed genes
A673
CHLA10
log Geneset Score
E
PDX sample UMAP
PDX-352
PDX-184
PDX-861
PDX-1058
PDX-856
F
r: -0.11
EWS-FLI1 activated genes
HOXD13 activated genes
PDX-1058
PDX-184
PDX-352
PDX-856
PDX-861
G
r: 0.49 (A673) r: 0.72 (CHLA10)
mesenchyme development (GO:0060485)
HOXD13 activated genes
H
r: 0.71
r: 0.47 (A673) r: 0.68 (CHLA10)
mesenchyme development (GO:0060485)
EWS-FLI1 repressed genes
I
r: 0.62
mesenchyme development (GO:0060485)
HOXD13 activated genes
J
r: 0.51
mesenchyme development (GO:0060485)
EWS-FLI1 repressed genes

**Figure S6. Single-cell expression of EWS-FLI1 and HOXD13 regulated gene sets**

Data were generated using CITE-seq. **A)** Low dimensional embedding, Uniform Manifold Approximation Projection (UMAP), of single-cell gene expression profiles of A673 and CHLA10 cells.

**B)** Ewing cells colored by HOXD13 activated gene set.

**C)** Ewing cells colored by Kinsey EWS-FLI1 activated gene set.

**D)** Ewing cells colored by Kinsey EWS-FLI1 repressed gene set.

**E)** Low dimensional embedding, Uniform Manifold Approximation Projection (UMAP), of single-cell gene expression profiles of 5 Ewing sarcoma PDX tumors (Aynaud et al. 2020).

**F)** Scatter plot of EWS-FLI1 activated genes and the HOXD13 activated genes by PDX.

**G)** Scatter plot of HOXD13 activated genes and the mesenchyme development gene set (GO:0060485) by cell line.

**H)** Scatter plot of EWS-FLI1 repressed genes and the mesenchyme development gene set (GO:0060485) by cell line.

**I)** Scatter plot of HOXD13 activated genes and the mesenchyme development gene set (GO:0060485) by PDX.

**J)** Scatter plot of EWS-FLI1 repressed genes and the mesenchyme development gene set (GO:0060485) by PDX.
